# Enhanced cultured diversity of the mouse gut microbiota enables custom-made synthetic communities

**DOI:** 10.1101/2022.03.07.483007

**Authors:** Afrizal Afrizal, Susan A. V. Jennings, Thomas C. A. Hitch, Thomas Riedel, Marijana Basic, Atscharah Panyot, Nicole Treichel, Fabian T. Hager, Ramona Brück, Erin Oi-Yan Wong, Alexandra von Strempel, Claudia Eberl, Eva M. Buhl, Birte Abt, André Bleich, René Tolba, William W. Navarre, Fabian Kiessling, Hans-Peter Horz, Natalia Torow, Vuk Cerovic, Bärbel Stecher, Till Strowig, Jörg Overmann, Thomas Clavel

## Abstract

Microbiome research is hampered by the fact that many bacteria are still unknown and by the lack of publicly available isolates. Fundamental and clinical research is in need of comprehensive and well-curated repositories of cultured bacteria from the intestine of mammalian hosts. In this work, we expanded the mouse intestinal bacterial collection (www.dsmz.de/miBC) to 212 strains, all publicly available and taxonomically described. This includes the study of strain-level diversity, small-sized bacteria, and the isolation and characterization of the first cultured members of one novel family, 10 novel genera, and 39 novel species. We demonstrate the value of this collection by performing two studies. First, metagenome-educated design allowed establishing custom synthetic communities (SYNs) that reflect different susceptibilities to DSS-induced colitis. Second, nine phylogenetically and functionally diverse species were used to amend the Oligo-Mouse Microbiota (OMM)12 model [Brugiroux et al. 2016 Nat Microbiol]. These strains compensated for differences observed between gnotobiotic OMM12 and specific pathogen-free (SPF) mice at multiple levels, including body composition and immune cell populations (*e*.*g*., T-cell subtypes) in the intestine and associated lymphoid tissues. Ready-to-use OMM stocks are available to the community for use in future studies. In conclusion, this work improves our knowledge of gut microbiota diversity in mice and enables functional studies via the modular use of isolates.

---

Omics technologies have been instrumental for exploring the diversity and functions within the gut microbiota, which include prokaryotes, fungi, and viruses, and for studying microbe-microbe and microbe-host interactions.^1^ A major challenge that remains is the large proportion of unknown microbial genes and corresponding taxa, which limits both molecular and experimental studies.^2,3^ The renewed interest in cultivation-based research on gut microbiomes helps address this issue, albeit primarily in the case of human gut bacteria.^4-8^ As enteric microbiomes are host species-specific and mice are important research models,^9-11^ we created the mouse intestinal bacterial collection (miBC) in 2016, making all strains publicly available.^12^ This included the first taxonomically described members of multiple novel genera, and the family *Muribaculaceae*, which has subsequently been reported by many.^13^ Since 2016, others have gathered bacterial isolates from the mouse intestine, albeit focusing on the *ob/ob* mouse model commonly used to study metabolic diseases (mouse gut microbial biobank; mGMB)^14^ or on functional differences between human and mouse gut microbiota (mouse gastrointestinal bacteria catalogue; MGBC).^15^ Despite these studies, many bacterial species remain either undescribed or unavailable in international culture repositories. Moreover, the utility of strains from these other collections for functional studies has not been demonstrated experimentally. Here we report new taxonomic and functional bacterial diversity from the mouse intestine, including the descriptions of 39 novel taxa, the study of strain-level diversity, and small-sized bacteria. We also present proof-of-concept experiments using miBC strains for modular functional investigation of microbe-host interactions. These experiments show a direct role of certain bacterial species in modulating immune responses and open avenues for study-specific synthetic communities (SYNs) of mouse gut bacteria.

## Results

### Expanding the cultured bacterial diversity from mouse gut microbiota

Diversity within the original collection released in 2016^12^ was doubled by including 112 bacterial strains, representing 73 fully-described species (reaching 141 species for the entire collection). This was achieved by obtaining isolates using different samples and culture conditions as specified in **Supplementary Table S1** and in the methods. The strains have been processed at international culture collections to guarantee long-term public availability. Their metadata and nucleotide sequences (near full-length 16S rRNA gene sequences and draft or closed genomes) can be accessed via the project repositories: www.dsmz.de/miBC and https://github.com/ClavelLab/miBC. The phylogenomy and occurrence in the mouse gut of new collection members are depicted in **Fig. 1**. Diversity was enriched across all bacterial phyla, including multiple strains of *Mucispirillum schaedleri* within the phylum Deferribacteres. The collection represents a total of six phyla, dominated by Firmicutes, and 35 families, dominated by *Lachnospiraceae* and *Lactobacillaceae*. Insights into novel bacterial diversity are presented in the next section. When several strains of the same species were obtained from mice of different origins, they were retained within the collection (**Supplementary Table S1**). In particular, due to the role of *Enterobacteriaceae* under dysbiotic conditions in multiple disease contexts,^16-19^ we included 20 *Escherichia coli* isolates with various origins, genomic and phenotypic features. All *E. coli* strains fermented lactose, a hallmark of this species compared with neighbouring members of the genus *Shigella*. In contrast, they varied in their ability to express flagella and their susceptibility to infection by known and newly isolated lytic phages^20^ (**Supplementary Fig. S1**). This toolbox will facilitate experiments to study community dynamics within SYNs at the strain level.

**Figure 1:**
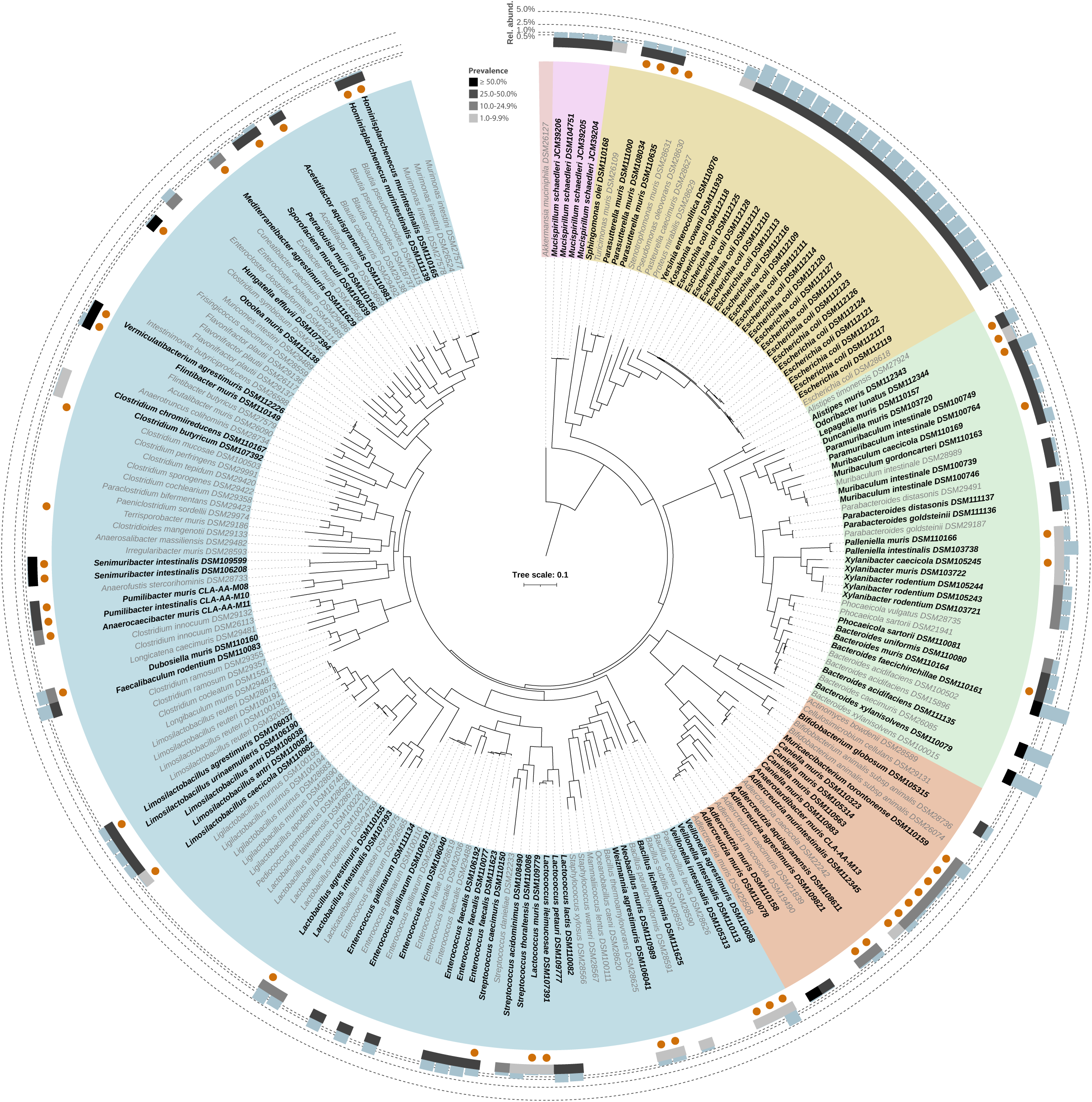
Phylogenomic tree of the mouse intestinal bacterial collection. The genomes used are listed in **Supplementary Table S1**. The tree was constructed using PhyloPhlAn v3.0.60^91^ and visualized and further processed in iTOL.^92^ For contextualization, isolates that were part of the first version of miBC^12^ are written in grey. Colours indicate phyla. Strains that are the first cultured members of novel taxa are indicated with orange dots surrounding their names. For all new isolates, the grey boxes and blue bars in the outer rings indicate the prevalence and mean relative abundance, respectively, of the corresponding 16S rRNA gene sequence in 11,485 amplicon datasets from mice.^21^

We then investigated how well the cultured isolates within miBC cover the mouse gut microbiota diversity as detected by sequencing. Analyses were performed in comparison with the two aforementioned resources of isolates recently published by others (mGMB and MGBC),^14,15^ with the limitation that MGBC does not provide full-length 16S rRNA gene sequences. We observed that 73 miBC isolates were shared with mGMB based on 16S rRNA genes at <98.7% sequence identity, yet miBC had almost twice as many isolates not accounted for by mGMB (134 *vs*. 77 sequences; representing 110 species) (**Fig. 2a**). A similar pattern was observed at the genome level (<95% ANI value), although the third collection MGBC contained an even greater number of unique isolates (n = 141) (**Fig. 2b**). Significant overlaps between the three collections were observed, accounting for approximately half of the genomes within each resource. The 101 genomes unique to miBC represented 89 species-clusters (>=95% ANI). This indicates that expanding culture collections, as done here, is helpful not only for increasing strain-level diversity of isolates across different countries, but also to provide unique bacterial diversity not yet captured by others.

**Figure 2:**
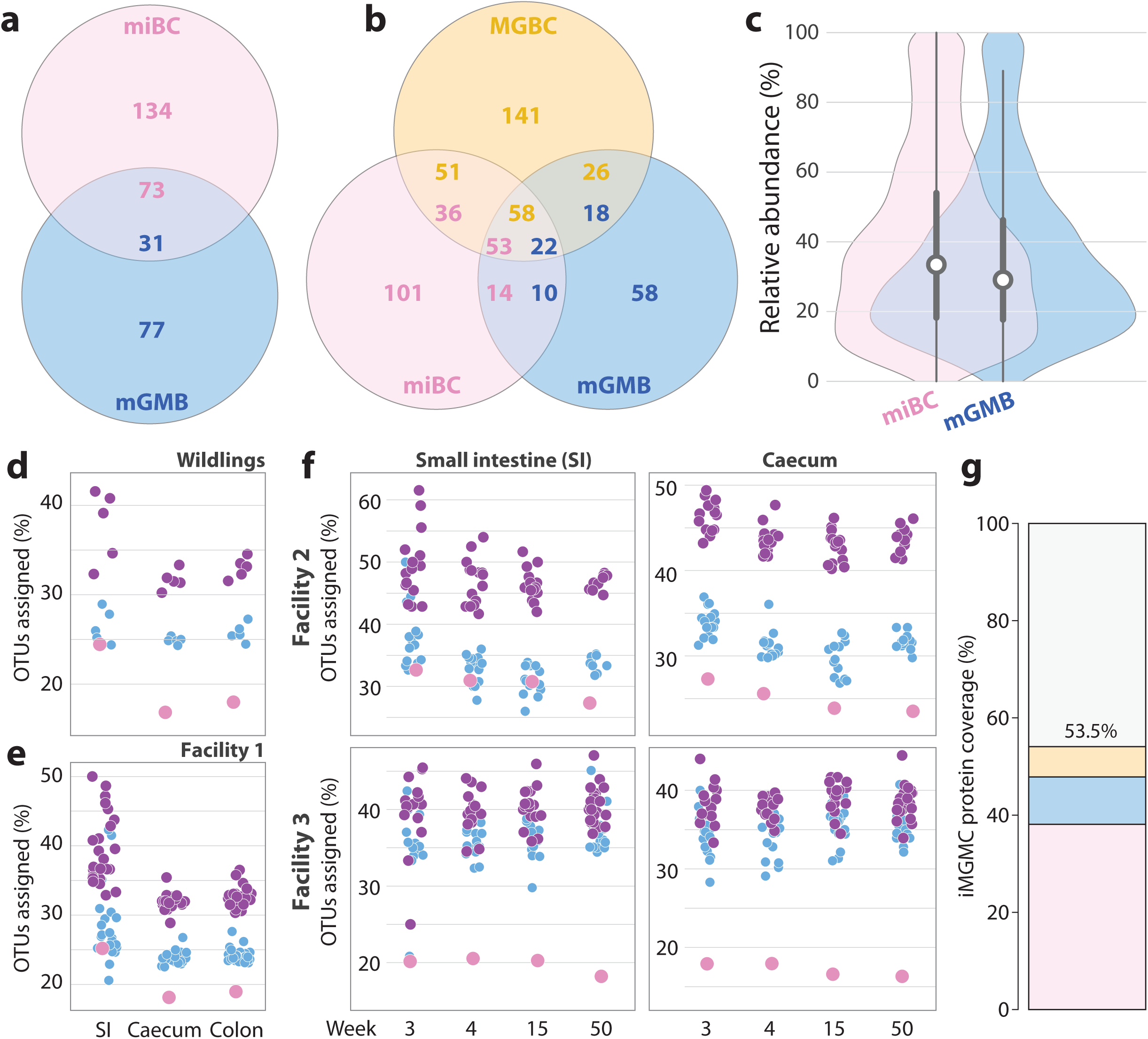
Comparison to other published collections of isolates and determination of cultured fractions. **(A)** Shared and unique diversity within miBC and mGMB^14^ based on 16S rRNA genes. Sequences with a pairwise identity >98.7% were considered to represent the same species. MGBC^15^ does not provide full-length 16S rRNA gene sequences and was not included in this analysis. **(B)** Shared and unique diversity within miBC, mGM, and MGBC^15^ based on genomes. ANI values >95% were used to define genomes representing the same species. **(C-F)** Cultured fractions (% of captured molecular species at the conservative threshold of 97 % sequence identity due to sequence size) of 16S rRNA gene amplicon data from mouse gut samples: **(C)** in the IMNGS database (n = 11,485),^21^ or **(D-F)** generated in the present study, including **(D)** caecum from wildling mice, **(E)** caecum from SPF mice in our own animal facility in Aachen (1), **(F)** various gut regions and different ages in two independent mouse facilities (2, TU Munich; 3, Medical School Hannover). Dots are colored as follows: pink, miBC; blue, mGMB; violet, mGMB plus unique diversity within miBC. **(G)** The percentage of proteins within the iMGMC gene catalog assignable to genomes within each collection was determined sequentially from miBC, mGMB and MGBC.

The cultured fraction of sequencing-based diversity was then assessed at three levels. First, using 11,485 amplicon datasets of mouse gut samples retrieved from IMNGS (>5,000 high-quality 16S rRNA gene sequences per sample),^21^ the median relative abundance accounted for by miBC was 36.2% at the genus and 27.0% at the species level (**Fig. 2c**). Compared with our original collection, this significantly improved coverage by 14.0% (genus) and 9.8% (species) (p<0.0001; Wilcoxon Rank-Sum). The coverage of the expanded miBC was also significantly better (p=0.002) than mGMB,^14^ which covered 35.3% at the genus and 24.4 % at the species level.

Second, we used 16S rRNA gene amplicon data from laboratory mice in different facilities and from wildling mice known to have a more diverse gut microbial ecosystem.^22^ The microbiota structure of laboratory mice depended on both the facility and gut region considered (ileum and caecum) (**Supplementary Text** and **Supplementary Fig. S2**). We observed substantial shifts in diversity and composition after weaning, without further consistent changes due to ageing (up to 50 weeks) compared with passive microbiota shifts within the given facility. The OTU coverage by miBC varied between facilities and was similar to mGMB in the small intestine (except in Facility 3), yet generally lower in the distal gut (**Fig. 2d-f**). This implies that several of the 134 isolates uniquely present in miBC represent taxa not captured by amplicon sequencing, either because they belong to sub-dominant populations or because some of them are generally missed by the method (*e*.*g*., DNA extraction efficiency; see next section on small-sized bacteria). Nevertheless, the miBC-unique cultured species clearly increased the fraction of amplicon sequences covered by sequences from the isolates within mGMB alone (**Fig. 2d-f**; violet dots).

Third, functional coverage was assessed at the metagenomic level using a recently published gene catalogue of the mouse gut.^23^ The expanded miBC collection covered 37.7% of all proteins in this dataset. When supplemented with the mGMB-and MGBC-derived genomes, a further 9.6% and 6.1% of proteins were accounted for, respectively (**Fig. 2g**). This means that miBC includes the majority of functions from bacterial strains cultured so far and that the three collections together cover over half (53.5%) of all functions detected by shotgun sequencing within the murine gut.

### Novel taxa and diversity of small-sized bacteria

The present cultivation work allowed to discover 39 novel bacterial taxa, which were described using Protologger,^24^ including taxonomic, ecological, and functional features based on near full-length 16S rRNA gene and genome sequences. Cell morphology was assessed by scanning electron microscopy (https://github.com/ClavelLab/miBC). These analyses led to the proposal of one novel family, 10 novel genera, and 39 novel species. Amongst them, the highest number of CAZymes was 410 in the genome of *Bacteroides muris*, suggesting that this species plays a role in carbohydrate degradation in the mouse gut. In contrast, the three novel *Adlercreutzia* species as well as *Anaerotardibacter muris* (all members of family *Eggerthellaceae*) had the lowest CAZymes repertoire (≤80 enzymes per genome). The pathway for sulfate assimilatory reduction to sulfide (EC:2.7.7.4, 2.7.1.25, 1.8.4.8, 1.8.1.2) was only detected in *Neobacillus muris* and *Weizmannia agrestimuris* (novel species and genus, respectively, within family *Bacillaceae*). Whilst the species *Odoribacter lunatus* (family *Odoribacteraceae*, phylum Bacteroidetes), which forms peculiar crescent-shaped cells, was absent from any of the 11,845 16S rRNA amplicon datasets analysed, *Alistipes muris, Otoolea muris*, and *Senimuribacter intestinalis* were highly prevalence in the mouse gut (>50% of samples positive for these species). Detailed information about all new bacteria obtained in this work is provided in **Supplementary Table S1** and in the protologues listed at the end of the methods section.

A specific protocol that we followed to successfully isolate novel bacteria was to pass gut suspensions through filters with a pore size of 0.45 µm to select for small-sized cells. This proved to be efficient in obtaining not only several strains of the species *Mucispirillum schaedleri*, as reported previously,^25^ but also three novel species distantly related to members of the family *Christensenellaceae*.^26^ According to their phylogenomy (**Fig. 3a)** and additional taxonomic analyses (see protologues), these isolates are proposed to be the first cultured members of a novel family, for which the name *Pumilibacteraceae* is proposed. Whilst certain isolates obtained via this filtration protocol grew indeed as small cells only, *e*.*g*., cocci with a diameter <0.5 µm in the case of strain CLA-AA-M08^T^ (**Fig. 3b**), others formed thin but long cells or displayed a more classical morphology, albeit with marked inter-cell heterogeneity, possibly explaining why some cells could pass the filter during preparation (see electron micrographs under https://github.com/ClavelLab/miBC). All three *Pumilibacteraceae* species and *Anaerotardibacter muris* (novel genus within family *Eggerthellaceae*) require relatively long incubation time to reach visible growth, are very sensitive to oxygen, and grow on agar medium only. A sufficient amount of genomic DNA could be obtained from *A. muris* only when additional enzymatic steps were included in the protocol, indicating that this species is difficult to lyse. Based on genome analysis, the three novel species within family *Pumilibacteraceae* were predicted to produce acetate, both from acetyl-CoA (EC:2.3.1.8, 2.7.2.1) and a combination of sulfide and L-serine (EC:2.3.1.30, 2.5.1.47). Moreover, they seem unable to utilise many carbohydrates, which co-occurred with a minimal CAZymes repertoire (<150 enzymes).

**Figure 3:**
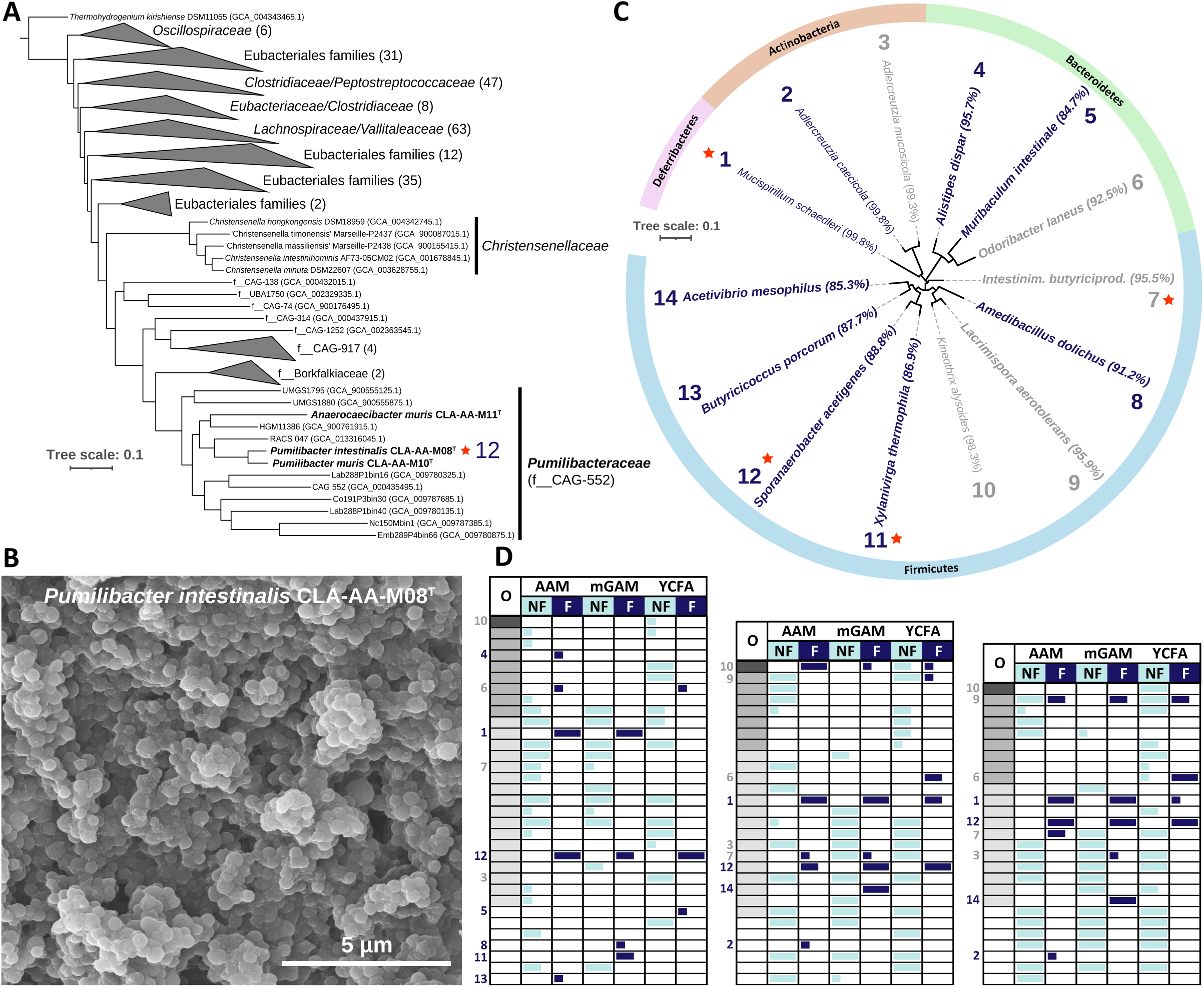
Diversity of small-sized bacteria in the mouse gut. **(A)** Phylogenomic tree of bacteria within the order *Eubacteriales*. The tree was constructed as in Figure 1. For clear visualisation, branches were collapsed whenever appropriate; in such cases, the number of species represented by the triangles are written in brackets after the corresponding family names. The genomes from isolates obtained in the present study (bold letters) are accessible via the project repository (see Data Availability section). Metagenome-assembled genomes were obtained from GTDB^93^; they are shown with their abbreviation from the database and corresponding accession number in brackets. **(B)** Scanning electron micrograph of strain CLA-AA-M08, for which the name *Pumilibacter intestinalis* within the novel family *Pumilibacteraceae* is proposed. Sample preparation is described in the methods section. **(C)** Phylogenetic tree based on the 16S rRNA gene amplicon sequences of dominant (>1 % relative abundance) operational taxonomic units (OTUs) obtained after culturing filtrates of mouse gut content. The OTU IDs were determined using EzBioCloud^61^ and are labelled using the closest relative with a valid name; the corresponding sequence identity is in brackets. Bold letters indicate OTUs considered to represent novel taxa at the conservative threshold of <97 % identity due to amplicons. Orange stars indicate the novel taxa first cultured and described in the present study. Blue letters indicate taxa exclusively found in the cultures from mouse gut filtrates and not in any other samples (original gut content or cultures thereof without pre-processing by filtration). **(D)** Presence of dominant OTUs (>1 % relative abundance) across the different types of samples and cultures. See methods section for detailed information. Three mouse caecal samples were cultured after filtration (F) or without (NF) in three different media (AAM, mGAMB, YCFA) in triplicates. The 16S rRNA gene amplicon profiles in these cultures were compared to that of the original samples (O). The grey gradient in these original samples indicate relative abundances as follows (from dark grey to white): >10%, 1-5%, 0.1-1%, <0.1%. Across the samples and culture media in this map, bars indicate the number of triplicate cultures positive for the corresponding OTU at >1 % relative abundance. Only those OTUs detected in at least one culture replicate across the entire set of samples are shown in this map. The complete dataset can be found in **Supplementary Table S2**. The bars in dark blue indicated OTUs that could be cultured from gut filtrates. These OTUs are numbered as in panel C, with dark blue numbers indicating those found exclusively in cultures from gut filtrates; those in grey letters were also found in cultures from unfiltered samples.

As a few intriguing isolates were obtained after 0.45 µm-filtration of gut content suspensions as presented above (**Supplementary Table S1**), we sought to characterise the diversity of such bacteria in a broader manner independent of the tedious handling and identification of single strains. Therefore, three freshly collected samples from laboratory mice were analysed by high-throughput 16S rRNA gene amplicon sequencing either as such or after filtration and cultivation on three different agar media, each in triplicate (see Methods section and data in **Supplementary Table S2**). Cultures from unfiltered caecal slurries served as controls. The diversity of taxa detected as dominant members of cultured communities (>1 % relative abundance) is shown in **Fig. 3c**. Out of the 14 molecular species spanning four phyla that were obtained from filtered material, 10 were considered to represent novel taxa (values <97 % in brackets and bold letters in the tree), of which three corresponded to the pure cultures mentioned above and are described in this work (orange stars). Moreover, nine of the 14 molecular species were exclusively present in the cultured communities obtained after filtration (see names and numbers in dark blue in **Fig. 3c** and **Fig. 3d**). These experiments demonstrate that easy-to-implement processing steps during sample preparation prior to cultivation allow the selection of specific taxa that would be otherwise too difficult to obtain directly from native communities. Whilst one of these small bacteria obtained as pure culture (no. 12, *Pumilibacter muris* CLA-AA-M08^T^ within the proposed novel family) was recovered in all media of all three samples tested and occurred at a relative abundance of ca. 0.3 % in the original sample *vs*. up to >90 % after filtration (**Fig. 3d**), others occurred in only one instance from filtered material (one replicate of one of the media for one given sample; blue bars). This shows that increasing the scale of such a work will be necessary in the future to capture an even broader range of novel mouse gut bacteria.

### Metagenome-based design of synthetic communities (SYNs) to study differential host responses

To demonstrate the value of a well-curated collection of mouse gut bacterial isolates to perform functional experiments, we first adopted a modular approach for synthetic community (SYN) design to generate consortia that mimic differential metagenomic functions. In this example, we generated SYNs associated with host susceptibility to DSS-induced colitis. For this purpose, the genomic cultured diversity in miBC and in shotgun metagenomes previously generated from mice of different origins, and characterized by varying disease severity after DSS treatment,^27^ were used as a foundation for data-driven SYN design using a modified version of our recently published bioinformatic workflow MiMiC (see methods).^28^ The binary (presence/absence) metagenomic profiles of protein families (Pfams) from the original faecal samples were clearly distinct in mice susceptible to DSS colitis (**Fig. 4a**). Whilst the two generated SYNs both consisted of species within the phyla Firmicutes and Bacteroidetes, their species composition differed markedly, with only two isolates being shared (**Fig. 4b**). This difference was less pronounced at the functional level, with 898 Pfams being unique and 3,985 shared between the two consortia, **Fig. 4c**. However, these unique functions were important enough to cause each consortium to better cover the respective samples they were derived from (**Fig. 4d**), especially in the case of mice resistant to DSS colitis where R-syn covered 4% additional functions than S-syn. When mapped to KEGG, these differences in Pfam coverage translated to a greater range of functional modules (231 *vs*. 220) and a greater metabolic capacity (metabolic pathways (map01100); 922 KOs *vs*. 865). This implies that the loss of commonly present functions (Pfams unique in metagenomes linked to the resistant phenotype) was partly responsible for the susceptibility to inflammation in this DSS model. Separation of the SYNs was also clearly observed on the multidimensional scaling plot and imitated the profiles of the original samples (**Fig. 4a**). SCFAs have long been known to impact gut health and both butyrate and propionate have been shown to improve resistance to DSS-induced colitis.^29,30^ In general, multiple pathways were observed to be less fragmented within the R-syn than S-syn. For instance, whilst both communities contained enzymes involved in the production of butyrate and propionate, more complete KEGG pathways were observed in the R-syn (propionate (map00640), 36 *vs*. 28; butyrate (map00650), 38 *vs*. 33). Propionate production could be followed from either succinate or glycerone phosphate to propanoyl-CoA in R-syn, which leads to three possible routes for propionate production, while both pathways for propanoyl-CoA production were incomplete in S-syn.

**Figure 4:**
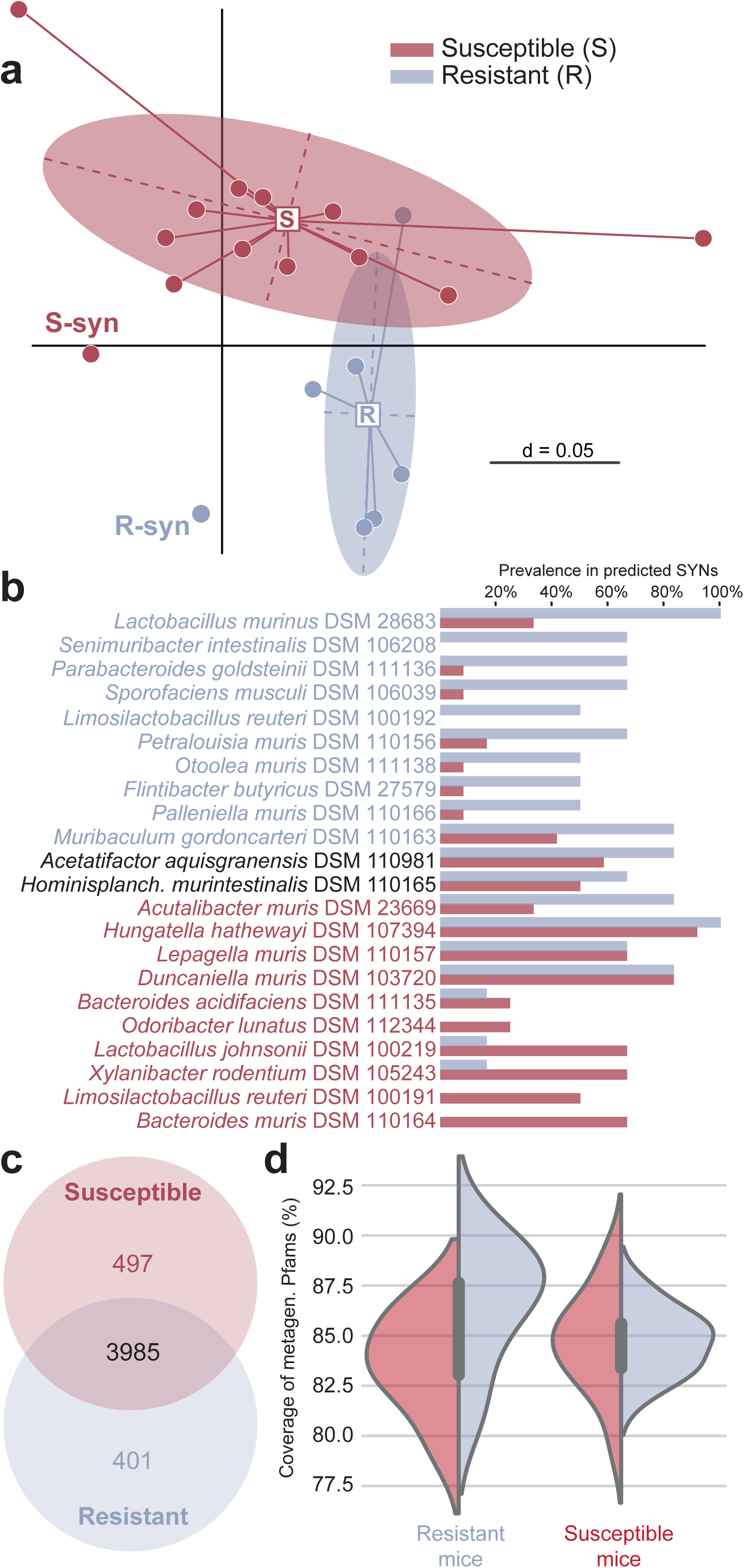
Modular design of synthetic communities (SYNs) to influence the severity of DSS-induced colitis in gnotobiotic mice. **(A)** Functional diversity of the original samples and the predicted SYNs as determined by multi-dimensional plotting of Jaccard distances based on binary protein family vectors calculated from shotgun metagenomic data. **(B)** List of bacterial strains included in the two SYNs (red, sensitive to colitis; blue, resistant). The two strains written in black letters were selected in both cases. The bars indicate how often a strain was selected within sample-specific SYNs in each group of mice (sensitive Vs. resistant). **(C)** Number of shared and unique Pfams between the two categories of SYNs. **(D)** Pfam-based functional coverage of the input metagenomic samples by the two SYNs.

### Colonization profiles of a new reference SYN

A few low-diversity mouse microbiota used as reference gut communities have been published to date,^31-33^ including the Oligo-Mouse Microbiota (OMM). The original OMM consists of 12 bacterial strains from the mouse intestine, herein termed OMM12.^31^ OMM12 has been used multiple times as described or complemented with one or more additional strains to study either microbe-host interactions or the ecosystem itself under controlled conditions, demonstrating the usefulness of such experimental models.^20,34-39^ However, due to the absence of important microbial functions in the OMM12, we selected additional phylogenetically and functionally diverse species from miBC to create the OMM19.1 model. We subsequently performed gnotobiotic experiments in two mouse facilities to validate colonization profiles and to study differential effects on the host. The selected strains and their features are presented in **Supplementary Fig. S3**. Ready-to-use strain mixtures of both OMM12 and 19.1 are publicly available for further use (www.dsmz.de/miBC).

Three sets of experiments were performed to test colonization by the OMM19.1 strains: (1) targeted colonization of germfree mice after weaning in gnotobiotic facility A (Aachen, Germany) to test engraftment in different gut regions; (2) colonization from birth using a breeding scenario in the same facility to test vertical transmission; (3) colonization after weaning in an independent gnotobiotic facility B (Hannover Medical School, Germany) to validate results. Bacterial composition was monitored by 16S rRNA gene amplicon sequencing with confirmation by qPCR for ileum and colon samples in colonization trial 1. All but one of the OMM19.1 species, *Flintibacter butyricus*, colonized the mice successfully, at varying relative abundances depending on gut regions (**Fig. 5a** and **Supplementary Fig. S4**). Stable colonization by *Extibacter muris* and *Escherichia coli* in this model agrees with previous findings.^34,37^ The relative abundance of *Bacteroides caecimuris*, a dominant member in the caecum and colon of OMM12 mice, was consistently reduced by colonization of the OMM19.1 strains, most likely due to the addition of *Parabacteroides goldsteinii* and *Xylanibacter rodentium*, two members of the same order (*Bacteroidales*) which were the most abundant OMM19.1 strains in the distal gut. In the small intestine, the dominance of *Akkermansia muciniphila* in OMM12 mice was apparently affected by colonization with *Ligilactobacillus murinus*. Interestingly, *Enterococcus faecalis* was not detected by both amplicon sequencing and qPCR in the intestine of OMM19.1, even though it was present in all three gut regions in OMM12 controls, suggesting that this species was affected by the added strains. Colonization by *M. schaedleri* in the colon of OMM19.1 mice (sporadically in the small intestine) was not seen in sequencing data but was confirmed by qPCR, albeit at low relative abundances (**Supplementary Fig. S4**). This agrees with the preferred habitat of this species being mucosa-associated areas.^40^ *Bifidobacterium animalis* was also detected in the colon of OMM12 mice by qPCR, but not in OMM19.1 counterparts.

**Figure 5:**
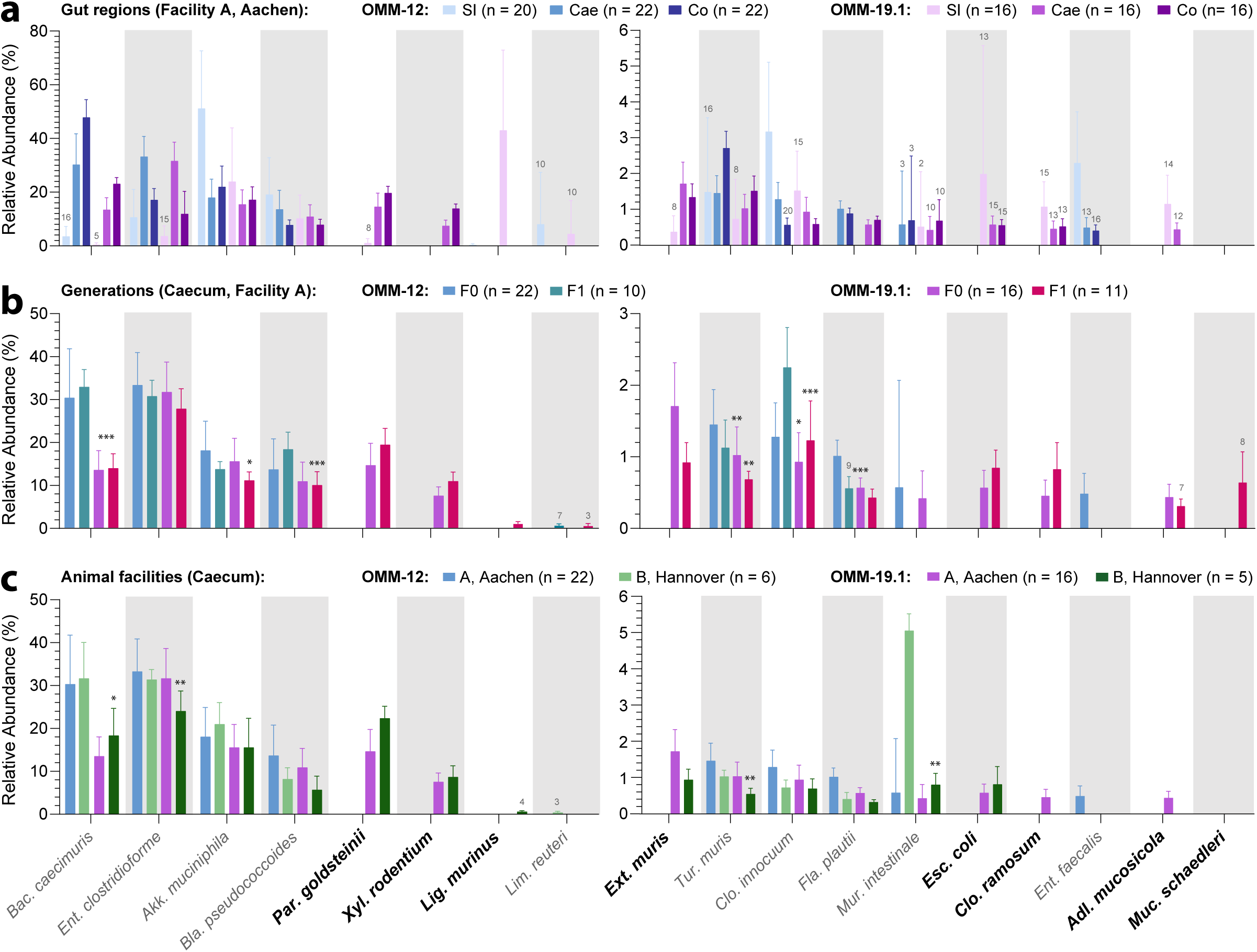
Colonization profiles in gnotobiotic mice inoculated with the original version of Oligo-Mouse Microbiota (OMM12) and its extended version OMM19.1. Detailed information on the strains added to the original OMM12 model is given is **Supplementary Figure S3**. These bacteria are written in bold letters below the x-axis. The number of mice included in each group are written in the figure **(A)** Bacterial composition in different gut regions in gnotobiotic facility A (RWTH Aachen) as obtained by high-throughput 16S rRNA gene amplicon sequencing. Samples from the caecum were also analysed by qPCR; these data can be seen in **Supplementary Figure S4**. All the bacteria detected in any of the experiments are consistently shown in the same order in all figure panels. For the sake of clarity, bacteria occurring at a relative abundance <1 % are shown in separate graphs and the y-axes have been optimized for visualisation of the values (right panels). Data in the caecum of mice in Facility A was used as a reference point in all figure panels (OMM12, blue bars; OMM19.1, violet bars. All values are shown as mean ± standard deviations. The total numbers of mice analysed in each experiment are indicated in brackets in the corresponding colour code legend. The numbers of mice positive for a species are shown in grey above the corresponding plot whenever inferior to the total number of mice. Abbreviations: SI, small intestine; Cae, caecum; Co, colon. **(B)** Bacterial composition in the caecum of mice from the F1 generation (Facility A). Detailed description as in A. **(C)** Bacterial composition in the caecum of mice from a second gnotobiotic facility (Facility B, Medical School Hannover). Detailed description as in A. P-values: * <0.05, ** <0.01, *** <0.001 (Mann-Whitney U-test; OMM12 *vs*. OMM19.1).

Vertical transmission of the OMM19.1 members was confirmed for 15 of the 16 strains detected in the caecum of F0-mice and mean relative abundances across OMM19.1 members were altogether stable (**Fig. 5b**). *Muribaculum intestinale*, which was previously shown to be sensitive to colonization protocols,^41^ could not be detected by amplicon sequencing after breeding; *E. faecalis* was also not detectable anymore in OMM12 mice of the F1 generation. In contrast *M. schaedleri* was present in F1-mice although detected only by qPCR in F0-controls. Colonization profiles in the caecum of mice from a second facility confirmed the presence of all dominant members of the OMM19.1 communities (**Fig. 5c**). The following species, detected at low relative abundances in the first facility, were absent in the second, although colonization in other gut regions was not tested and qPCR was not performed: *Flavonifractor plautii, Clostridium ramosum, E. faecalis, M. schaedleri*. In summary, whilst it is expected that colonization profiles in future experiments may vary between gut regions and facilities (*e*.*g*., differences in diet and other environmental factors) and depend on the method used for detection, providing standardized stocks as a starting point for colonization and using validated colonisation protocols^41^ reduce the risk of variations. Moreover, such stocks delivered robust profiles for dominant members of the communities (>1 % relative abundance) in our experiments.

### Differential effects on the host

To compare effects of the different types of microbial communities on the host (OMM12 and OMM19.1 *vs*. germfree (GF) and specific pathogen-free (SPF) controls), mice colonized after weaning in facility A were phenotyped via body imaging and immune cell profiling in the intestinal lamina propria (LP) and gut-associated lymphoid tissues (GALT) by flow cytometry. For many of the parameters, OMM19.1 mice showed an intermediate state between OMM12 and SPF controls, even though results did not reach statistical significance for several single parameters due to inter-individual variabilities (**Fig. 6**). In contrast to an expected decrease in caecum weight due to colonization (**Fig. 6a**), total body weight and fat content were not different between groups (**Fig. 6a** and **6b**). However, interesting findings included increased heart and lung volume, as well as an increase in femur density (but not length) (**Fig. 6b**). In terms of immune readouts (**Fig. 6c** and **Supplementary Fig. S5**), most notable changes were observed in T cell subtypes and IgA+ plasma cells in various LP and GALT compartments. Whilst the overall proportion of CD4+ T cells did not differ between the groups, phenotypic composition of these cells was altered by the microbiota. RORγt+ CD4+ Th17 cells were nearly absent in GF mice but their prevalence increased with complexity of the microbiota in both LP and GALT. The fraction of Foxp3+ Tregs in the SI and mesenteric lymph nodes (MLNs) did not change with colonisation status but was increased in the colonic LP of colonised mice. Notably, within the Foxp3+ Treg population, RORγt-expressing Tregs were increased in OMM-19.1 mice in all compartments compared with GF and OMM-12 controls. Similarly, there was an overall increase in the frequency of IgA+ plasma cells in both intestinal LP compartments, with OMM-19.1 mice showing intermediate values between the OMM-12 and SPF groups. Few differences were observed in the proportion of other innate and adaptive immune cell populations (**Supplementary Fig. S5**).

**Figure 6:**
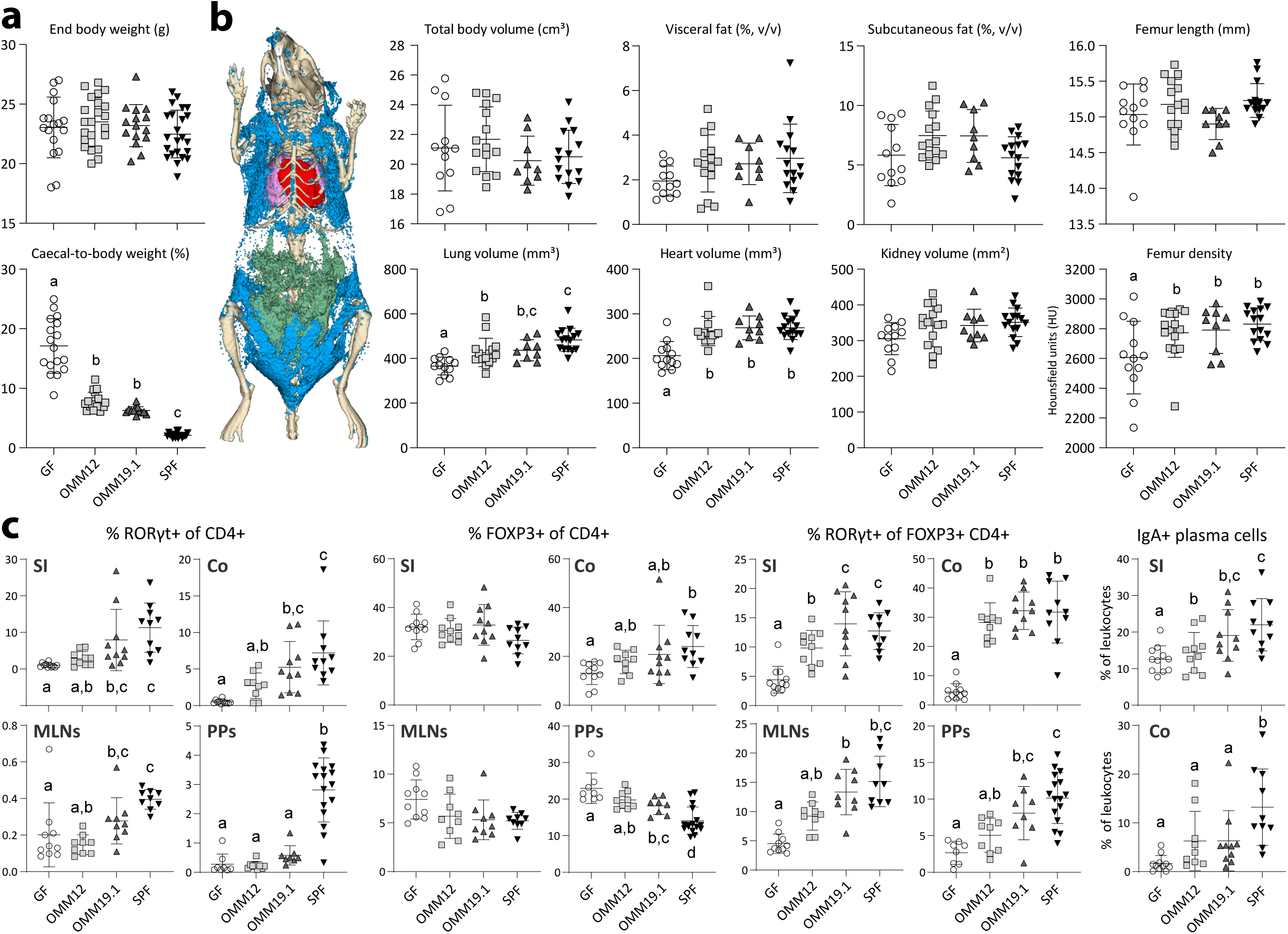
Differential effects of OMMs on the mouse physiology. Corresponding colonization profiles are shown in Fig. 4a. **(A)** Body and caecum weight measured at culling. The total number of mice in each group was: GF, n = 17; OMM12, n = 22; OMM19.1, n = 15; SPF, n = 23). **(B)** Body imaging data obtained as described in the method section (GF, n = 12; OMM12, n = 15; OMM19.1, n = 9; SPF, n = 15). **(C)** Immune phenotyping by flow cytometry. All leukocytes were initially gated as live CD45+ cells. CD4+ T cells were identified as TCRβ+ CD4+ and further subdivided into RORγt+ FoxP3-(Th17), FoxP3+ (Treg), and FoxP3+ RORγt+ subsets, as indicated. IgA+ plasma cells were identified as IgA+ B220-Ly6c+. The parent gate is indicated in the individual graphs. Other data can be seen in **Supplementary Fig. S5**. Different letters indicate values that are statistically significant between groups (Kruskall-Wallis test followed by Mann-Whitney U-test for pairwise comparisons). Numbers of mice were: (i) small intestine (SI) and colon (Co); GF, n = 11; OMM12, n = 9; OMM19.1, n = 10; SPF, n = 10; (ii) mesenteric lymph nodes (MLNs); GF, n = 10; OMM12, n = 9; OMM19.1, n = 9; SPF, n = 9; (iii) Peyer’s patches (PP); GF, n = 8; OMM12, n = 9; OMM19.1, n = 8; SPF, n = 16).

Taken together, this work provides access to strains and mixed consortia allowing for targeted colonization studies in gnotobiotic mice. Body composition and immune parameters demonstrated that implementing the OMM12 model with additional strains and their functions contributed to inducing host responses closer to conventionally colonized mice.

## Discussion

Mouse models are widely used in fundamental and clinical research. It is thus important to characterize in detail the factors that modulate their physiology, such as the gut microbiota. Despite work in the last five years,^12,14,15^ the diversity of yet-uncultured bacteria from the mouse intestine is still high, which hinders further advances in the field. Establishing state-of-the-art collections of mouse gut bacteria is a tedious endeavour due to the high-quality standards that are difficult to comply with and followed by very few. The final resource must be well curated, and the isolates must be made publicly available at the time of publication and, in the best case, fully characterized taxonomically. The present work aims towards these goals. Moreover, it provides insights into the contribution of isolates to pathophysiology of the host.

The expanded range of bacterial diversity within miBC makes new strains available to perform experimental studies more easily. For instance, *Mucispirillum schaedleri* was already included in altered versions of the Schaedler Flora,^33^ but access of this species has been problematic since then. The new strains provided here will help to further elucidate its ecology (*e*.*g*., enrichment in the intestine of rodents), lifestyle, and role in inflammation and resistance to infections.^42^ Compared with other published resources,^14,15^ we aim to provide detailed information on novel taxa and eventually validate their names to generate added value for the community. Our collection includes multiple novel species within important bacterial groups in the mouse gut, such as the *Muribaculaceae, Coriobacteriales*, or *Clostridiales*.^2,13^ Whilst this taxonomic distribution agrees overall with the diversity of isolates provided by others,^14,15^ each collection brings unique diversity to light. The present work reports full-description of 39 novel taxa, including one new family represented by small-sized bacteria.

There are only a few examples of synthetic communities (SYNs) from the mouse intestine that have been established and used since the 1960s.^31-33^ Such models can be implemented in two manners: (I) as reference communities to perform gnotobiotic studies under controlled and reproducible conditions;^32,38^ or (II) as modular systems to test the effects of specific strains added to the community.^34,37^ By amending the original OMM12 model via the addition of phylogenetically and functionally complementary strains and by making the corresponding strains mixtures and all single miBC strains available, the work presented here facilitates both types of models aforementioned. Previous work has highlighted the link between the gut microbiota and growth of the host, including bone-related parameters, especially under malnutrition.^43,44^ We found that increasing bacterial diversity of the OMM model to 19 strains shifted body composition towards the phenotype of conventionally colonized mice, including femur density and the size of several organs. A recently published community of 15 mouse isolates (GM15), which also included four miBC strains, was shown to increase femur length to values observed in OMM12 and SOPF/SPF controls.^32^ However, this community is not publicly available. Previous work pointed at the role of lactobacilli in maintaining growth of infant mice during chronic undernutrition.^44^ Whilst GM15 and OMM19.1 have two species within family *Lactobacillaceae* in common (*Limosilactobacillus reuteri* and *Ligilactobacillus murinus*),^45^ additional work will be needed to dissect the role of single OMM19.1 members in the phenotypes observed.

The bacterial strains used to establish the OMM19.1 model influenced immune cell populations in GALT, especially RORγt+ T cells and IgA+ plasma cells. It is sound to ask whether these effects are due to the increased diversity *per se* or to specific functions of the added strains. Some bacteria, particularly those associated with the intestinal epithelium, were reported to effectively induce adaptive immune responses, notably IgA production and RORγt+ T cell priming.^46,47^ Moreover, microbial colonisation is known to stimulate the maturation of germinal centres in GALT, resulting in the accumulation of IgA somatic mutations that influence microbiota reactivity.^36,48^ Recent studies highlighted the importance of secondary bile acids produced by gut bacteria in regulating the maturation and functions of Treg and Th17 populations.^49-51^ *Extibacter muris* is such a bacterium within OMM19.1 capable of producing secondary bile acids by 7*α*-dehydroxylation.^37^ Another recent study reported high induction of RORγt+ Tregs by the species *Clostridium ramosum*, which is also a member of OMM19.1.^52^ The fraction of IgA+ plasma cells was very low in the colon of GF mice and expanded substantially due to colonisation. The role of these and other bacteria^53^ in conferring the differential effects observed between OMM12 and 19.1 at the level of immune cells will be worth investigating in the future.

Variations in the gut microbiota of mouse models, *e*.*g*., due to the origin of mice, has been shown to markedly influence immune responses and the susceptibility of mice to DSS-induced colitis.^27,54^ Approaches used so far to design the composition of SYNs has primarily been taxa-centric, *i*.*e*., based on expert knowledge of the diversity and functions of isolates that are easy to culture. In a yet unpublished work, phenotype variations in the DSS-induced colitis model were investigated at a large-scale, emphasizing again the importance of gut microbes in this model.^55^ This study followed also a taxa-centric approach to identify novel species within the genera *Duncaniella*^13^ and *Alistipes*^56^ that play a role in disease onset. We here propose a function-based approach that, independent of taxonomic boundaries and based on the functional landscape within host-specific collections of isolates, provides SYN compositions that reflect dysbiotic conditions captured by shotgun sequencing. The biological relevance of such SYNs designed using individual metagenomic data remains to be tested in future studies. Whilst the relatively low diversity within SYNs is a limiting factor when compared to stool-derived *in vitro* communities,^57^ working with a controlled system is an important advantage. Moreover, such approaches based on custom-made SYNs adapted to the needs of specific studies will gain in power proportionally to further expansion of isolate collections.

In conclusion, the resource presented here provides multiple novel insights into the mouse gut microbiota that will hopefully facilitate the work by others on microbe-host interactions in health and disease. Next steps include further technical developments (both wet lab and *in silico*)^24,58^ to enhance the throughput and optimize the in-to-output ratio of cultivation approaches, especially regarding rapid and precise identification of isolates. Large-scale cultivation studies^5^ still report a low depth of cultivation per individual sample, and yet-unpublished work on the development of high-complexity SYNs^59^ is based on isolates of various origins. Further efforts are thus required to reach the goal of personalized microbiome-based research and applications using isolates.

## Methods

### Samples and culture media for bacterial isolation

Samples were collected from mice euthanized for scientific procedures in accordance with the German Animal Protection Law (TierSchG). The internal animal care and use committee (IACUC) at the University Hospital of RWTH Aachen approved the collection of gut content from donor mice not subjected to any experimental treatment (internal approval no. 70018A4). Gut content was also collected in the context of studies otherwise ethically approved by the federal authority (Landesamt für Natur, Umwelt und Verbraucherschutz, North Rhine Westfalia, LANUV; approval no. 81-02.04.2020.A131 and 81-02.04.2019.A065). Particular attention was paid by the experimenters during dissection and sampling to reduce the risk of potential contaminations by bacteria from the environment (*e*.*g*., use of sterile materials only, disinfected dissection set, thorough disinfection of the dissection area and the mice prior to dissection). Gut contents were collected into 2ml Eppendorf tubes immediately after culling and brought into an anaerobic workstation (MBraun, Garching, Germany) with an atmosphere consisting of 4.7 % H_2_ and 6 % CO_2_ in N_2_ and a partial pressure of oxygen <0.1 ppm. The list of culture media used in this study and their compositions are provided in **Supplementary Table S1**.

### Bacterial isolation and characterization

Sterile agar media were placed in the anaerobic workstation at least 24 hours before use. Fresh gut contents were re-suspended (1:10 wt/vol) in anaerobic, reduced (0.05% L-cysteine, 0.02% DTT) phosphate-buffered saline (PBS). The suspensions were then serially diluted in PBS down to 10^−6^. Each dilution was plated onto the agar media and incubated at 37 °C for up to 7 days. Single colonies were picked and re-streaked three times onto fresh agar plates to guarantee purity of the isolates. The strains were first identified using a MALDI-Biotyper (Bruker Daltonik), following the manufacturer’s instructions. For isolates not identifiable at the species-level by MALDI (score <1.7) or identified as species not yet contained in the original collection,^12^ the 16S rRNA gene was sequenced. Genes were amplified by PCR using primer 27f and 1492r.^60^ PCR products were purified and sent for Sanger sequencing at Eurofin Genomics (Ebersberg, Germany) or Microsynth Seqlab (Göttingen, Germany) using primer 27f, 1492r, 338r, and 785r. The raw sequences were first checked and modified manually with help of the electropherograms prior to building contigs to obtain near full-length 16S rRNA gene sequences. The most closely related species with a valid name were identified using Ezbiocloud.^61^ A cut-off value of 98.7% was used as a first layer of delineation between known and novel species.^62^

For phage infection assays, *E. coli* strains were incubated in BHI medium overnight at 37 °C and 100 µl of biomass was plated onto LB agar plates. After drying, 2.5 µl of phage lysate dilution series (in BHI medium down to 10^−7^) were pipetted onto the plate, each dilution in triplicate. After overnight incubation at 37 °C, productive lysis by the phages was observed visually by the appearance of individual plaques within the spotting zone at appropriate phage dilutions. For the assays with phage Mt1B1_P3, Mt1B1_P10 and Mt1B1_P17,^20^ the bacteria were streaked on LB agar plates from frozen stocks. After overnight incubation at 37°C, single colonies were resuspended in 90µl LB Medium of which 20µl were immediately streaked onto EBU (Evans blue, uranine) agar plates. After drying, 5µl of the phage lysates were spotted in duplicate and incubated overnight at 37°C. Bacterial lysis was observed visually by the appearance of plaques and colour change to a darker green of the agar plate around the spot. All *E. coli* strains were also further analysed using EnteroPluri-Test (Liofilchem®) to test for substrate utilisation. Single colonies from freshly grown strains on BHI agar were used for inoculation using sterile needles. The tests were visually assessed after incubation at 37 °C for 24 h according to the manufacturer’s instructions.

### Strain processing at the DSMZ

After shipment as live cultures at room temperature or cryo-stocks on dry ice, strains were cultured and quality checked following standard operating procedures at the Leibniz Institute DSMZ using the strain-specific conditions specified on the website (www.dsmz.de/miBC). Purity was confirmed by re-streaking whenever possible, visual observation of colony morphologies, and microscopic observation of cells. Isolates were assigned unique collection numbers and they were kept either as cryo-stocks in capillaries stored in liquid nitrogen or in lyophilized form using glass ampoules stored at 4 °C for long-term storage. The Oligo-Mouse Microbiota (OMM) strain mixtures (OMM10-basis; extension set 1.2; extension set 2.9) were prepared in an anaerobic workstation by mixing equal volumes of the corresponding strains freshly cultured separately under appropriate anaerobic conditions. The final stocks contained 12% glycerol as a cryo-protectant and were stored at -80 °C in crimp closed glass vials (Macherey-Nagel, ref. no. 70201HP and 70239).

### Flagellin assays

The Flagellin Bioactivity Assay was modified from a method described previously.^63^ HEK-BlueTM-hTLR5 cells were used, which were generated by co-transfection of the human TLR5 gene and an inducible SEAP (secreted embryonic alkaline phosphotase) reporter gene into HEK293 cells. The SEAP gene is placed under the control of the IFN-β minimal promoter fused to five NF-κB and AP-1-binding sites. Stimulation of the TLR5 receptor by ligands, such as flagellin, activates NF-κB and AP-1, which induce the production of SEAP. Activity is determined using QUANTI-BlueTM (Invivogen), with a change from pink to purple-blue colour indicating a positive reaction and thus TLR5 induction. Cells resuspended in maintenance medium (per litre of DMEM without L-glutamine: 100 mL heat-inactivated FCS, 1 mmol L-glutamine, 5 mL Pen/Strep, 200 µL normocin, 300 µL blasticidin, 100 µL zeocin) were pipetted into wells of a 96-well plate at a density of ca. 2.5 × 10^4^ cells per well. The *E. coli* strains used in these experiments are listed in **Supplementary Table S1** and presented in **Supplementary Fig. S1**. Suspensions of freshly grown *E. coli* strains in BHI medium (OD_600_, 0.5) were diluted 100-fold and 20 µl were added to the HEK-cells in triplicates. FLA-ST, standard flagellin from *Salmonella enterica* Typhimurium (Invivogen), was used to generate a standard curve (10-200 ng/ml). BHI medium without bacteria was used as a negative control. The plates were incubated at 37 °C under an atmosphere containing 5% CO2 for 21 h and then centrifuged (100 x g, 4 °C, 5 min). Supernatants (20 µl) were added to 180 µl of QUANTI-BlueTM (Invivogen) and incubated for 45 minutes. SEAP activity was measured at 630 nm using a microplate reader. The assay war repeated tree times for each strain.

### Genome sequencing and analysis

The biomass of freshly grown isolates revived from frozen glycerol stocks was collected from liquid media or agar plates. The DNA was isolated using a modified version of the method by Godon et al. (1997).^64^ Cells were lysed by bead-beating in the presence of DNAse inhibitor and detergent, then purified on NucleoSpin gDNA Clean-up columns (Macherey-Nagel, Germany). For isolates that were hard to lyse (*e*.*g*., CLA-AA-M13), additional enzymatic steps with lysozyme (Carl Roth, ref. 8259.1; 3 mg/L; 37 °C, 30 min) and proteinaseK (Carl Roth, ref. 7528.1; 500 mg/L; 50 °C, 1-2 h) was added prior to bead-beating. DNA integrity was checked by gel electrophoresis and concentration was measured using a Qubit fluorometer (Thermo Fischer Scientific, USA).

DNA libraries were prepared with the NEBNext Ultra II FS DNA Library Prep Kit for Illumina (NEB, USA) according to the manufacturer’s protocol using ∼300 ng of DNA. The time used for enzymatic shearing to ca. 200 bp was 30 minutes. The PCR enrichment of adaptor-ligated DNA was conducted with five cycles and NEBNext Multiplex Oligos for Illumina (NEB, USA) for paired-end barcoding. For size selection and clean-up of adaptor-ligated DNA, AMPure beads (Beckman Coulter, USA) were used. Quality check (Bioanalyzer System, Agilent Technologies, USA) and DNA quantification (Quantus, Promega, USA) of resulting libraries were conducted at the IZKF Core Facility (UKA, RWTH Aachen University), as was the sequencing on a NextSeq500 (Illumina, USA) with a NextSeq500 Mid Output Kit v2.5 (300 Cycles).

Raw reads were quality-filtered and adapters as well as phiX reads were removed using Trimmomatic v0.39^65^ and bbduk.^66^ Assemblies were obtained using SPAdes v3.13.1.^67^ Any contigs shorter than 1000nt were removed before quality check using CheckM.^68^ Genomes were deemed to be of high quality when >95% complete and containing ≤5% contamination. The N50, longest contig, shortest contig, total number of contigs and coverage were calculated using the ‘Assembly_stats.py’ code available at: https://github.com/thh32/Assembly-stats. Comparison to both mGMG^14^ and MGBC^15^ was done using all publicly available genomes at the time of analysis (October 2021). This consisted of 108 genomes for the mGMB collection, each representing a different species, and 276 for the MMGC collection, representing 132 species. Comparisons between 16S rRNA gene sequences was conducted using the code within Protologger^24^ with matches determined at 98.7% similarity. Genome comparison was conducted using FastANI using a match threshold of 95%.^69^

### Ageing mouse dataset

To accurately determine the cultured fraction of gut microbiota from laboratory mice (specific pathogen-free, SPF; C57BL/6 wildtype mice), the content of the small intestine and caecum from animals in two facilities were sampled at different ages: 3 weeks, 4 weeks, 15 weeks, and 50 weeks (Facility 2, Munich) or 45 weeks (Facility 3, Hannover). To avoid cage and litter effects, several litters and cages were sampled at each time-point. Moreover, to account for natural microbiota drifts within each facility over time, additional 10-week-old mice were used as controls and sampled at the earliest and latest time point. Routine microbiological monitoring allowed excluding infections by common murine pathogens.^70^ These experiments did not include any interventions on the mice, which were housed in controlled environments with water and standard chow diets provided *ad libitum*. All procedures were in accordance with the German Animal Welfare Legislation. For experiments in Facility 2, the breeding and sampling of wildtype animals for scientific purposes was according to Paragraph 4, Section 3, and did not require specific approval due to the absence of intervention. Use of the animals was documented in the yearly animal records sent to the authorities. Breeding and housing of the mice in the facility fulfilled all legal requirements according to Paragraph 11, Section 1, Sentence 1. For experiments in Facility 3, the procedures were approved by the local Institutional Animal Care and Research Advisory Committee and covered by the permission of the local veterinary authority (reference no. 2015/78).

### Mouse experiments with Oligo-Mouse Microbiota (OMM)

#### Experiments in gnotobiotic facility A (University Hospital of RWTH Aachen, Germany)

All experiments were performed under Ethical Approval LANUV no. 81-02.04.2019.A065 in accordance with EU regulation 2010/63/EU. All mice used were bred in germfree (GF) isolators (NKPisotec, Flexible film isolator type 2D) under sterile conditions. To obtain specific-pathogen free (SPF) mice with the same genetic background (C57BL/6N) as their GF counterparts, mice were taken from the isolator and housed with SPF mice, allowing passive colonisation with a complex microbiota. The first generation of conventionalised mice after breeding were taken for use in this work. The initial main cohort of mice were fed *ad libitum* using autoclaved (134 °C, 20 min) fortified standard chow (ssniff V1534-300) and given autoclaved water (pH 7), with F1 generation mice fed on irradiated fortified chow instead (ssniff V1124-927). GF mice were removed from breeding isolators at 5 weeks of age and housed in HEPA-filtered bioexclusion isocages (Techniplast ISO30P). In both isolators and isocages, Tek-Fresh bedding (ENVIGO) was used. Mice were housed in single sex cages in the same room. Room temperature was kept between 21-24 °C and 25-40% humidity on a 12h:12h day:night cycle. Faecal samples were taken before starting experiments to confirm the GF status via microscopic observation after Gram-staining and plating on both anaerobic and aerobic agar plates.

To assess effects of the OMM19.1 consortium (n = 16 mice) on the host, it was compared to OMM12 (n =22), GF (n = 20) and SPF (n = 23) controls. Each gnotobiotic group was created using age-matched GF mice, including animals from several litters and cages to account for potential confounding effects. The OMM-stocks (prepared as described above) were introduced orally by gavage (50 µl per mouse), followed by 100 µl rectally. A fresh aliquot was used for each cage, as recommended previously.^41^ The second dose was given after 72hrs. SPF controls (also age-matched) were taken from the conventionalised C57BL/6N sister line. OMM19.1 and -12 mice were also bred under gnotobiotic conditions to assess colonization profiles after vertical transmission. All mice were culled at the age of 13 weeks, *i*.*e*., after 8 weeks of colonization for gavaged mice. Gut content from the small intestine, caecum, and colon was collected for bacterial composition analysis by 16S rRNA gene amplicon sequencing and quantitative PCR (qPCR). Body imaging was carried out as described below. Small intestinal and colonic tissues as well as mesenteric lymph nodes (MLNs) and Peyer’s patches (PPs) were collected during dissection and processed immediately for immune phenotyping by flow cytometry, as described below.

#### Experiments in Gnotobiotic Facility B (Institute for Laboratory Animal Science, Hannover Medical School, Germany)

This study was conducted according to the German animal protection law and European Directive 2010/63/EU. All experiments were approved by the Local Institutional Animal Care and Research Advisory committee and permitted by the Lower Saxony State Office for Consumer Protection and Food Safety (LAVES; file no. 18A367 and 2018/188). GF male and female C57BL/6JZtm mice were obtained from the Central Animal Facility (Hannover Medical School, Hannover, Germany). They were maintained in plastic film isolators (Metall+Plastik GmbH, Radolfzell-Stahringen, Germany) in a controlled environment and twelve-hour light/dark cycles. Hygiene monitoring according to standard operating procedures^70,71^ confirmed that the mice were free of contaminants or infection with common murine pathogens throughout the experiment. GF mice (n = 3 males and 3 females per group) were removed from the breeding isolator at the age of five weeks and colonized with the corresponding OMM stocks, SPF microbiota, or left untreated (GF controls). Colonization occurred twice, 3 days apart, using 50 µl orally and 100 µl rectally of fresh OMM stocks. One OMM aliquot was used for each cage. The SPF group was colonized by following the same procedure but using freshly prepared caecal slurries from SPF, C57BL/6J mice. After colonization, mice were kept in airtight cages with positive pressure (IsoCage P, Tecniplast Deutschland GmbH, Bavaria, Germany) and received pelleted 50 kGy gamma-irradiated feed (Sniff) and autoclaved water *ad libitum*. Each group was created using age-matched GF mice, including several litters and cages to account for potential confounding effects. Mice were culled via CO_2_ inhalation followed by exsanguination at 13 weeks of age (8 weeks post inoculation) and caecal samples were collected to measure bacterial colonisation by amplicon sequencing.

### Body imaging

On the day of sampling, mice were weighed, anesthetised with 2% isoflurane in air, and imaged using a micro-computed tomography device (U-CT, MILabs B.V.). Ultra-focus fast scan mode with a resolution of 0.16 mm x 0.16 mm x 0.16 mm, tube voltage of 65 kV, tube current of 0.13 mA and a scan time of 27 s was used for fat analysis. Ultra-focus normal scan mode was used to segment organs and bones. The resolution was the same but tube voltage, tube current and scan time were 55 kV, 0.17 mA and 3 min 42s, respectively. All µCT Scans were reconstructed by MILabs Auto Rec 1.6, organs and fat were segmented using Imalytics Preclinical 2.1.9.11 (Gremse-IT).^72^

### Immune cell phenotyping by flow cytometry

Intestinal tissues were cut longitudinally and washed in HBSS/3% FCS (Hank’s Balanced Salt Solution/Foetal Calf Solution) to remove any gut content. Peyer’s patches (PPs) were removed for separate analysis and excess fat was cut away. Small intestine (SI) and large intestine (LI) were cut into 5-mm sections and shaken vigorously in HBSS 2 mM EDTA to remove epithelial cells. Samples were then incubated at 37 °C with shaking for 20 min, after which they were filtered through a 50µm Nitex mesh (Sefar) and the supernatant discarded. The remaining sample was washed with HBSS. These steps were repeated again using fresh HBSS 2mM EDTA. A final rinse using HBSS was performed to remove trace amounts of EDTA and filtered through the Nitex mesh. Depending on the tissue being studied, one of the following methods was used: (i) for the isolation of leukocytes from the lamina propria (LP) of SI, the tissue was placed in 15ml RPMI containing 1mg/ml collagenase VIII (Sigma, C2139-1G); (ii) for colonic tissue (LI), a mix of enzymes was used (collagenase V, Sigma, C9263-1G, 0.85 mg/ml; Collagenase D, Roche, 11088882001, 1.25 mg/ml; dispase, Gibco, 17105-041, 1 mg/ml; DNase, Roche, 101104159001, 30µg/ml). Tissues were then incubated at 37 °C with shaking for at least 15 min and manually shaken every 5 minutes until complete digestion. Tubes containing the tissue were then placed on ice, filtered through a 100-µm cell strainer, and centrifuged (400 x g, 6 min).

For mesenteric lymph nodes (MLNs), after removal of any remaining fat, they were placed in a 1.5-ml sample tube containing 500 µl RPMI (without FCS) and cut into small pieces. 500 µl FCS free RPMI containing 2 mg/ml Collagenase D were added to each tube (end concentration 1 mg/ml). Tissue was incubated at 37°C for 45 min under constant shaking. The cells were strained through a 100 μm cells strainer into a 50 ml tube, washed with PBS/3% FCS and centrifuged (400 x g, 6 min, 4°C).

PPs were excised from the SI and digested (37 °C, 45 min) using 100 µg/ml liberase TH/DNase (Roche) in RPMI containing 5 % FCS. Mononuclear phagocytes were enriched by MACS using CD11c magnetic beads (Miltenyi) according to the manufacturer’s protocol. The CD11c negative fraction was used for subsequent analysis of B and T cells.

Flow cytometry staining and analysis was performed at the IZKF Flow Cytometry Core Facility of the RWTH University Hospital. For surface staining, single cell suspensions from SI, colon, MLNs and PPs were centrifuged (400 x g, 6 min) and resuspended in PBS/3% FCS containing a mix of fluorescently labelled antibodies for identification of different cell populations. Antibodies against mouse Ly6c (HK1.4), MHCII (M5/114.15.2), Ly6G (1A8), CD11b (M1/70), B220 (RA3-6B2), CD19 (6D5), CD64 (X54-5/7.1), CD103 (2E7), CD11c (N418), CD4 (RM4-5), CD8α (53-6.7), TCRβ (H57-597), TCRγδ (GL3) were purchased from BioLegend, IgA (mA-6E1) from eBioscience and CD45 (30-F11) from BD. Cells were stained with 7-AAD for viability (Biolegend).

For intracellular staining, cell suspensions were stained with the Zombie NIR fixable viability dye (Biolegend) according to manufacturer’s instructions and incubated for 20 minutes at 4 °C. The cells were washed in PBS/3% FCS, centrifuged (400 x g, 6 min) and resuspended in PBS/3% FCS containing antibodies for surface staining as described above. The cells were stained for 45 minutes at 4°C in the dark, then washed in PBS/3% FCS, centrifuged (400 x g, 6 min) and the cell pellets were resuspended in 1X fixation buffer (TF staining kit, eBioscience) overnight at 4°C in the dark. Fixed cells were centrifuged at 400 x g for 6 minutes and the pellet resuspended in 1 x permeabilisation buffer (TF staining kit, eBioscience). The cells were centrifuged (400 x g, 6 min) and then resuspended in 1 x permeabilisation buffer containing the antibodies for intranuclear staining: Foxp3 (FJK-16s) and RORγt (B2D) from eBioscience. The cells were stained at room temperature for 1h, then washed in PBS/3% FCS, centrifuged (400 x g, 6 min) and finally resuspended in PBS/3% FCS for flow cytometry.

The cells were acquired on a BD LSRFortessa flow cytometer (BD) and analysed using the FlowJo analysis software (BD).

### High-throughput 16S rRNA gene amplicon analysis

Samples were processed and analysed as described previously.^58^ In brief, metagenomic DNA was purified on columns (Macherey-Nagel) after mechanical lysis by bead-beating. The V3-V4 regions of 16S rRNA genes were amplified (25 cycles), purified using AMPure XP magnetic beads (Beckman-Coulter, Germany) and sequenced in paired-end mode using the v3 chemistry (600 cycles) on an Illumina MiSeq according to the manufacturer’s instructions. The platform was semi-automated (Biomek4000 pipetting robot, Beckman Coulter, Germany) to increase reproducibility and the workflow systematically included two negative controls (a DNA-extraction control, *i*.*e*., sample-free DNA-stabilization solution, and a PCR blank, *i*.*e*., PCR-grade water as template) for each 46 samples sequenced. Raw sequencing reads were processed using IMNGS (www.imngs.org),^21^ a platform based on UPARSE.^73^ A sequence identity threshold of 97% was used for clustering sequences into operational taxonomic units (OTUs). Unless otherwise stated, only OTUs that occurred at a relative abundance ≥0.25% in at least one sample were kept for further processing.^74^ OTUs were taxonomically classified using SILVA (Pruesse et al., 2012). Further data processing (diversity and composition analyses) was done in R using Rhea.^12^

To determine cultured fractions, the 16S rRNA gene sequences of isolates were matched to OTUs using blastn (E-value <1e−25, 97% identity, 80% query coverage). Large-scale ecological analysis of the mouse gut was conducted using 11,485 datasets downloaded from the IMNGS database.^21^ 16S rRNA amplicon samples containing ≥5000 sequences and labelled as ‘mouse gut’ (n = 11,485) were used. The 16S rRNA gene sequences of all isolates were compared to IMNGS-derived OTU sequences using blastn (E-value <1e−25, 97% identity, 80% query coverage).

### 16S rRNA-gene targeted quantitative PCR (qPCR)

16S rRNA gene-targeted primers and probes for the nine new strains within OMM19.1 were designed as described before,^31^ except those for *Mucispirillum schaedleri, Escherichia coli*, and *Extibacter muris*, which were published elsewhere.^20,34,37^ Sequences were: *Adlercreutzia mucosicola*, f-5’GCTTCGGCCGGGAAT, r-5’GGCAGGTTGGTCACGTGTTA, Hex-CAGTGGCGAACGGGTGA-BHQ1; *Clostridium ramosum*, f-5’GCGAACGGGTGAGTAATACATAAGT, r-5’GCGGTCTTAGCTATCGTTTCCA, Fam-ACCTGCCCTAGACAGG-BHQ1; *Xylanibacter rodentium*, f-5’AAGCGTGCCGTGAAATGTC, r-5’CGCACTCAAGGACTCCAGTTC, Hex-CTCAACCTTGACACTGC-BHQ1; *Parabacteroides goldsteinii*, f-5’CGCGTATGCAACCTACCTATCA, r-5’ACCCCTGTTTTATGCGGTATTAGTC, Fam-AATAACCCGGCGAAAGT-BHQ1; *Flintibacter butyricus*, f-5’TAGGCGGGAAAGCAAGTCA, r-5’CAAATGCAGGCCACAGGTT, Fam-ATGTGAAAACCATGGGC-BHQ1; *Ligilactobacillus murinus*, f-5’TCGGATCGTAAAACCCTGTTG, r-5’ACCGTCGAAACGTGAACAGTT; Hex-TAGAGAAGAAAGTGCGTGAGAG-BHQ1. For absolute quantification of 16S rRNA copy numbers, standard curves using 10-fold dilution series (1-10^6^ copies/µl) of linearized plasmids containing the target sequence were generated using 6 replicates. qPCR assays and specificity testing were performed as described previously (Brugiroux et al, 2016).

### Shotgun metagenome analysis

For comparison to the metagenomic gene catalogue, the protein sequences from the genome of each isolate were extracted using Prodigal (v2.6.3) using default options. These sequences were then annotated against the protein sequences within iMGMC^23^ using DIAMOND blastp (v2.0.8.146),^75^ with a minimal bit-score of 100.

For the prediction of synthetic communities using MiMiC,^28^ host reads were removed from the metagenomic samples using BBmap based on the methods defined in the iMGMC pipeline.^23^ The filtered metagenomic reads were assembled using Megahit (v1.2.9) with default options.^76^ Protein sequences were then extracted from each assembly using Prodigal (v2.6.3), with the ‘-p meta’ flag.^77^ Proteins were then annotated against the Pfam database (v32)^78^ using HMMscan,^79^ filtered using the gathering threshold option (--cut_ga). For each sample, the annotation was converted into a binary presence/absence vector file. The genome of each isolate was also annotated against the Pfam database and used to generate a binary presence/absence vector file for comparison against the metagenomic samples. MiMiC scoring was modified to include weighting (score modifier = 0.0005) for each Pfam present in >50% of samples within a group. Secondly, weighting (score modifier = 0.0012) was applied to Pfams that occurred significantly more frequently (Fischer exact, p-value < 0.05) within either of the groups. An initial round of sample-wise consortia selection was conducted for each group of mice (sensitive *vs*. resistant to DSS-induced colitis). The isolates selected within at least three samples within a group were included in a reduced list of isolates used for a second group-wise selection. In this second level of selection, each group-specific reduced list of isolates was used to generate a list of all potential 12-member consortia. For each group, the vector of each consortium was compared to the vectors of all individual samples, each providing a MiMiC score as described above. The consortium with the highest consortia-wide MiMiC score across each group of samples was selected as being most representative of that group.

### Taxonomic description of novel bacteria

The general scheme followed here to describe novel taxa was as described in our recent work.^58^ In brief, draft genomes were generated for the strains supposed to represent novel taxa due to a 16S rRNA gene sequence identity <98.7% to any bacteria with a valid names.^61,80^ This was followed by taxonomic, ecological, and functional analyses using Protologger (www.protologger.de).^24^ All raw output files of these analyses are available in the project data repository: https://github.com/ClavelLab/miBC. For each isolate, taxonomy was assigned using the following thresholds: <98.7% (as indication for a novel species), <94.5% (novel genus), and <86.5% (novel family) based on 16S rRNA gene sequence similarities;^81^ ANI values <95% and genome-based differences in G+C content of DNA >1%^82^ to separate species; POCP values <50% for distinct genera.^83^ Phylogenomic trees were also considered to make decisions on genus-and family-level delineation. Manual POCP analysis, Genome-to-Genome Distance Calculator 3.0^84^ with a cut-off of 70% for species level, and ANI Calculator^61^ were also performed for the delineation of certain species. Scanning electron micrographs of all the isolates representing novel taxa are available online (https://github.com/ClavelLab/miBC). In addition to the 36 novel taxa described here in the expanded version of miBC, the protologues below also include the description of three isolates from the original collection,^12^ for which genomes have now been generated, revealing their status of novel species.

### Description of *Acetatifactor aquisgranensis* sp. nov

*Acetatifactor aquisgranensis* (a.quis.gra.nen’sis. M.L. masc. adj. *aquisgranensis*, pertaining to Aachen (Germany), where the bacterium was isolated). The isolate has the highest 16S rRNA gene sequence similarity to *Acetatifactor muris* (89.17%). Despite this relatively low value that may indicate a novel genus, GTDB-Tk classified the genome as an unknown species within the genus *Acetatifactor*. The closest relative based on genome tree is *A. muris*, which shares a POCP value of 59.45%, above the genus delineation threshold. As the ANI and GGDC value of the isolate to *A. muris* are 81.00% and 29.10%, respectively, the isolate is proposed to represent a novel species within the genus *Acetatifactor*. Cells are generally straight to slightly bent rods (1-2µm in length) when grown in BHI medium for 3-7 days at 37°C under anaerobic conditions. In total, 404 CAZymes were identified within the genome. The ability to utilise starch and cellulose as carbon source was predicted. KEGG analysis identified pathways for the production of acetate from acetyl-CoA (EC:2.3.1.8, 2.7.2.1), propionate from propanoyl-CoA (EC:2.3.1.8, 2.7.2.1), L-cysteine and acetate from sulfide and L-serine (EC:2.3.1.30, 2.5.1.47), L-glutamate from ammonia via L-glutamine (EC:6.3.1.2, 1.4.1.-), and folate (vitamin B9) from 7,8-dihydrofolate (EC:1.5.1.3). No antibiotic resistance genes were identified. The 16S rRNA gene sequence of the species was most prevalent in the mouse gut (47.6% of 1,000 samples positive, at an average relative abundance of 0.23%), followed by human vagina (1.9%). The type strain is **CLA-AA-M01 (=DSM110981**^**T**^). Its G+C content of genomic DNA is 51.2 mol%. It was isolated from the caecal content of an SPF mouse.

### Description of *Adlercreutzia agrestimuris* sp. nov

*Adlercreutzia agrestimuris* (a.gres.ti.mu’ris. L. masc. adj. *agrestis*, wild; L. masc. or fem. n. *mus*, a mouse; N.L. gen. n. *agrestimuris*, of a wild mouse). The next relatives based on 16S rRNA gene sequence analysis was *Enteroscipio rubneri* (92.10% identity), followed by members of the genus *Adlercreutzia*. GTDB-Tk assigned the genome to an unknown genus within family *Eggerthellaceae*. Phylogenomic analysis confirmed that this isolate belongs to family *Eggerthellaceae*, forming a separate branch within a cluster of species from the genus *Adlercreutzia*. POCP values >50% were observed for multiple genera within family *Eggerthellaceae*: *Senegalimassilia anaerobia* (family *Coriobacteriaceae* in LPSN), 58.5%; *Adlercreutzia caecimuris*, 58.4%; *Eggerthella lenta*, 54.3%; *E. rubneri*, 58.*1%; Slackia piriformis*, 57.0%; *Berryella intestinalis*, 55.8%. However, the highest POCP value was 61.7% to *Adlercreutzia equolifaciens*, the type species of this genus. The isolate also shared POCP values >50% to other strains from this study proposed to represent additional novel *Adlercreutzia* species: *Adlercreutzia murintestinalis* (58.7%) and *Adlercreutzia aquisgranensis* (64.0%). None of the close species with a valid name and these other isolates shared ANI and GGDC values above the corresponding species delineation thresholds. Altogether, with the current state of isolates and genomes available from members of the family *Eggerthellaceae*, the taxonomic placement of novel taxa is ambiguous due to conflicting data. Despite relatively low 16S rRNA gene sequence identities and the GTDB-Tk assignment of this isolate, we propose to create a novel species within the genus *Adlercreutzia*, and not a novel genus, to avoid generating more confusion. This decision was primarily based on highest POCP value to *Adlercreutzia equolifaciens* (the type species of this genus) and phylogenomic placement of the isolate. The taxonomy of genera and species within family *Eggerthellaceae* will have to be consolidated in the near future when a higher number of isolates and genomes are available. Cells are short rods (ca. 0.6-1.2 µm in length) when grown in WCA medium under anaerobic conditions for up to 5 days. In total, 75 CAZymes were identified within the genome. No genes for carbon source utilisation were found. KEGG analysis identified pathways for the production of acetate from acetyl-CoA (EC:2.3.1.8, 2.7.2.1), propionate from propanoyl-CoA (EC:2.3.1.8, 2.7.2.1), L-cysteine and acetate from sulfide and L-serine (EC:2.3.1.30, 2.5.1.47), and L-glutamate from ammonia via L-glutamine (EC:6.3.1.2, 1.4.1.-). No antibiotic resistance genes were detected. The 16S rRNA gene sequence of the species was most prevalent in the mouse gut (7.1% of 1,000 samples positive) at low relative abundance. The type strain is **CLA-SR-6**^**T**^ **(=DSM 109821**^**T**^). Its G+C content of genomic DNA is 48.2%. It was isolated from the gut content of a wild mouse.

### Description of *Adlercreutzia aquisgranensis* sp. nov

*Adlercreutzia aquisgranensis* (a.quis.gra.nen’sis. M.L. fem. adj. *aquisgranensis*, pertaining to Aachen (Germany), where the bacterium was isolated). Based on 16S rRNA gene sequence comparisons, the isolate was most closely related to *Adlercreutzia muris* (94.82%). However, GTDB-Tk assigned the genome to an unknown genus within family *Eggerthellaceae*. Phylogenomic analysis confirmed that this isolate belongs to family *Eggerthellaceae*, as it forms a separate branch within a cluster of species from the genus *Adlercreutzia*. The isolate shares POCP value >50% to species from multiple genera within family *Eggerthellaceae*, albeit with highest value of 60.5% to *Adlercreutzia caecicola* and 59.5% to *Adlercreutzia equolifaciens*, the type species of this genus. The isolate also shared POCP values >50% to other strains from this study proposed to represent additional *Adlercreutzia* novel species: *Adlercreutzia murintestinalis* (53.6%) and *Adlercreutzia agrestimuris* (64.0%). Based on (i) the 16S rRNA genes sequence identity above the genus delineation threshold (94.5%), (ii) high POCP values to several *Adlercreutzia* spp., and (iii) phylogenomic analysis, we think it is sound to place this isolate within the genus *Adlercreutzia*. None of the close species with a valid name and the other isolates aforementioned shared ANI and GGDC values above the corresponding species delineation thresholds, confirming the status of novel species. Cells are rods (0.6-1.2 µm in length) when grown in WCA medium under anaerobic conditions for up to 5 days. In total, 73 CAZymes were identified within the genome. No genes for carbon source utilisation were found. KEGG analysis identified pathways for the production of acetate from acetyl-CoA (EC:2.3.1.8, 2.7.2.1), propionate from propanoyl-CoA (EC:2.3.1.8, 2.7.2.1), and L-glutamate from ammonia via L-glutamine (EC:6.3.1.2, 1.4.1.-). No antibiotic resistance genes were detected. The 16S rRNA gene sequence of the species was most prevalent in the mouse gut (14.5% of 1,000 samples positive) at low relative abundance. The type strain is **CLA-RA-2**^**T**^ **(=DSM 108611**^**T**^). Its G+C content of genomic DNA is 64.4%. It was isolated from the gut content of a wild mouse.

### Description of *Adlercreutzia murintestinalis* sp. nov

*Adlercreutzia murintestinalis* (mur.in.tes.ti.na’lis. L. mas. or fem. n. *mus*, a mouse; N.L. masc. adj. *intestinalis*, intestinal; N.L. fem. adj. *murintestinalis*, of the mouse intestine). The closest relatives based on 16S rRNA gene sequence similarities are *Adlercreutzia equolifaciens* subsp. *equolifaciens* (92.21%), followed by *Adlercreutzia equolifaciens* subsp. c*elatus* (91.64%), and *Adlercreutzia caecicola* (91.30%). GTDB-Tk assigned the isolate to an unknown genus within family *Eggerthellaceae*. Phylogenomic analysis confirmed that this isolate belongs to family *Eggerthellaceae*, forming a separate branch within a cluster of species from the genus *Adlercreutzia*. POCP values above the genus delineation threshold (50%) were obtained against members of multiple genera within family *Eggerthellaceae*, including: *Adlercreutzia* (*A. equolifaciens*, 54.12% (highest); *Adlercreutzia caecimuris*, 51.74%; *Adlercreutzia mucosicola*, 53.27%), *Gordonibacter* (*Gordonibacter urolithinfaciens*, 51.99%), and *Senegalimassilia* (*Senegalimassili*a *anaerobia*, family *Coriobacteriaceae* in LPSN, 50.31%). The isolate also shared POCP values >50% to other strains from this study proposed to represent additional novel *Adlercreutzia* species: *Adlercreutzia agrestimuris* (58.7%) and *Adlercreutzia aquisgranensis* (53.6%). However, none of the close species with a valid name and these other isolates shared ANI and GGDC values above the respective species delineation thresholds. Altogether, with the current state of isolates and genomes available from members of the family *Eggerthellaceae*, the taxonomic placement of novel taxa is ambiguous due to conflicting data. Despite relatively low 16S rRNA gene sequence identities and the GTDB-Tk assignment of this isolate, we propose to create a novel species within the genus *Adlercreutzia*, and not a novel genus, to avoid generating more confusion. This decision was primarily based on highest POCP value to *Adlercreutzia equolifaciens* (the type species of this genus) and phylogenomic placement of the isolate. The taxonomy of genera and species within family *Eggerthellaceae* will have to be consolidated in the future when a higher number of isolates and genomes are available. The genome contained only 80 CAZymes and no carbohydrate utilisation pathways were identified. KEGG analysis identified pathways for acetate production from acetyl-CoA (EC:2.3.1.8, 2.7.2.1) and propionate production from propanoyl-CoA (EC:2.3.1.8, 2.7.2.1). Ecological analysis suggested that the species is most prevalent within amplicon datasets from the mouse gut (5.6%). The type strain is **CLA-AA-M17** ^**T**^ **(=DSM 112345**^**T**^). Its G+C content of genomic DNA is 59.3 mol%. It was isolated from the gut content of an SPF mouse.

### Description of *Alistipes muris* sp. nov

*Alistipes muris* (mu’ris L. gen. n. *muris* of a mouse). This isolate showed highest 16S rRNA gene sequence similarities to species within the genus *Alistipes* (*Alistipes dispar*, 95.79%; *Alistipes timonensis*, 95.59%; *Alistipes putredinis*, 95.30%). GTDB-Tk identified the genome as species “Alistipes sp002428825”. The genus assignment was supported by POCP analysis (60.22% to *A. timonensis*) and by the placement of the isolate amongst *Alistipes* species in the genomic tree, including the type species of this genus, *A. putredinis*. None of the closest relatives shared an ANI value above 95% with the genome of this isolate, confirming its status as a novel species. Cells were rods, mostly 1.0-2.5 µm in length when grown on YCFA Agar for 3-10 days at 37°C under anaerobic conditions. The total number of CAZyme indentified in the genome was 140. No genes related to carbon source utilisation were found. KEGG analysis identified pathways for the production of acetate from acetyl-CoA (EC:2.3.1.8, 2.7.2.1), propionate from propanoyl-CoA (EC:2.3.1.8, 2.7.2.1), and L-glutamate from ammonia via L-glutamine (EC:6.3.1.2, 1.4.1.-). The detection of major facilitator superfamily (MFS) antibiotic efflux pump (ARO:0010002) may indicate resistance to antibiotics. The 16S rRNA gene sequence of the species was most prevalent in the mouse gut (51.1% of 1,000 samples positive, at an average relative abundance of 1.98%), followed by pig (3.6%) and human gut (3.0%). The type strain is **CLA-AA-M12 (=DSM112343**^**T**^). Its G+C content of genomic DNA is 59.2 mol%. It was isolated from filtered (0.45 µm) gut content of an SPF, Fsp27^-/-^ mouse.

### Description of *Anaerocaecibacter* gen. nov

*Anaerocaecibacter* (An.ae.ro.cae’ci.bac.ter. Gr. pref. *an-*, not; Gr. masc. n. *aer*, air; L. neut. n. *caecum*, caecum; N.L. masc. n. *bacter*, rod; N.L. masc. n. *Anaerocaecibacter*, an anaerobic rod from caecum). The closest relatives based on 16S rRNA gene similarity are *Xylanivirga thermophila* (83.57%), *Christensenella hongkongensis* (83.57%), *Caldicoprobacter guelmensis* (83.41%), and *Caldicoprobacter faecalis* (83.29%). POCP values to all close relatives were below 30% and GTDB-Tk placement assigned the type species to an unknown genus within ‘f_CAG-552’. Separation from the other proposed species *Pumilibacter muris* and *Pumilibacter intestinalis* within the propose novel family *Pumilibacteraceae* (see protologue below) was confirmed by phylogenomic placement, which showed they were distinct from each other, and by the POCP value of 45.9% between the type species of each genus. The type species is *Anaerocaecibacter muris*.

### Description of *Anaerocaecibacter muris* sp. Nov

*Anaerocaecibacter muris* (mu’ris L. gen. n. *muris* of a mouse). Cells are rods (length: 1.2-2.7 µm, diameter ca. 0,5µm) when grown on YCFA or mGAM Blood agar under anaerobic conditions for 2-4 weeks. In total, 93 CAZymes were identified within the genome of the type strain and only starch was predicted to be utilised as a carbon source. KEGG analysis identified pathways for the production of acetate from acetyl-CoA (EC:2.3.1.8, 2.7.2.1), propionate from propanoyl-CoA (EC:2.3.1.8, 2.7.2.1), L-cysteine and acetate from sulfide and L-serine (EC:2.3.1.30, 2.5.1.47) and L-glutamate from ammonia via L-glutamine (EC:6.3.1.2, 1.4.1.-). Antibiotic resistance was predicted based on the detection of tetracycline-resistant ribosomal protection protein. Ecological analysis suggested that the species is most prevalent within amplicon datasets from the mouse gut (16.3%). The type strain is **CLA-AA-M11**^**T**^. Its G+C content of genomic DNA is 50.3 mol%. It was isolated from a filtered (0.45 µm) faecal suspension of an SPF, Fsp27^-/-^ mouse.

### Description of *Anaerotardibacter* gen. nov

*Anaerotardibacter* (An.ae.ro.tar.di.bac’ter. Gr. pref. *an-*, not; Gr. masc. n. *aer*, air; L. masc. adj. *tardus*, slow; N.L. masc. n. *bacter*, a rod; N.L. masc. n. *Anaerotardibacter*, slow growing anaerobic rod, pertaining to the slow growing nature of the bacterium). The isolate shares highest 16S rRNA gene sequence similarity to *Eggerthella sinensis* (89.61%). GTDB-Tk assigned the genome to an unknown genus ‘CAG-1427’ within the family *Eggerthellaceae*. The phylogenomic tree analysis placed the isolate within the cluster containing multiple genera from the *Eggerthellaceae*, with the closest relative being *Denitrobacterium detoxificans*. However, the POCP value to this species was 47.6%, while the highest value was to *Senegalimassilia anaerobia* (50.3%). These analyses support the creation of a novel genus to accommodate this isolate. The type species is *Anaerotardibacter muris*.

### Description of *Anaerotardibacter muris* sp. Nov

*Anaerotardibacter muris* (mu’ris L. gen. n. *muris* of a mouse). The species shares all features of the genus. Cells are short rods (0.6-1.2 µm in length) when grown on YCFA or mGAM Blood (5%) agar under anaerobic conditions for 1-3 weeks, as the bacterium is a slow grower. Very low DNA amount could be extracted when no enzymatic lysis was added during extraction. The total number of CAZymes identified in the genome was 73. No genes for carbon source utilisation were predicted. KEGG analysis identified pathways for the production of acetate from acetyl-CoA (EC:2.3.1.8, 2.7.2.1) and propionate from propanoyl-CoA (EC:2.3.1.8, 2.7.2.1). No antibiotic resistance genes were detected. Ecological analysis suggested that the species is most prevalent within amplicon datasets from the human gut (17.3% of 1,000 samples positive), followed by wastewater (14.5%), and mouse gut (7.1%). The type strain is **CLA-AA-M13**^**T**^. Its G+C content of genomic DNA is 54.2 mol% It was isolated from the gut content of an SPF, Fsp27^-/-^ mouse.

### Description of *Bacteroides muris* sp. nov

*Bacteroides muris* (mu’ris L. gen. n. *muris* of a mouse). According to 16S rRNA gene sequence analysis, this bacterium was most closely related to *Bacteroides* spp. (max. 96.69% to *Bacteroides uniformis*). GTDB-Tk identified the genome as ‘Bacteroides sp002491635’. Assignment to the genus *Bacteroides* was also supported by POCP analysis, with highest value of 67.38% to *B. uniformis*, and 54.0% to *B. fragilis* (type species). In the genome tree, the isolate formed a cluster with *B. uniformis* and *B. rodentium*. However, the corresponding ANI and GDGC values were below the species delineation thresholds (92.08%/48.20% and 90.92%/43.10%, respectively), justifying the proposal to create a novel species within the genus *Bacteroides*. The bacterium grows well on Columbia blood agar (5% sheep blood) within 24 hours at 37°C under aerobic conditions. The total number of CAZymes identified within the genome was 410. The ability to utilise starch and cellulose as carbon source was identified. KEGG analysis revealed pathways for the production of acetate from acetyl-CoA (EC:2.3.1.8, 2.7.2.1), propionate from propanoyl-CoA (EC:2.3.1.8, 2.7.2.1), L-cysteine and acetate from sulfide and L-serine (EC:2.3.1.30, 2.5.1.47), L-glutamate from ammonia via L-glutamine (EC:6.3.1.2, 1.4.1.-), folate (vitamin B9) from 7,8-dihydrofolate (EC:1.5.1.3), and riboflavin (vitamin B2) from GTP (EC:3.5.4.25, 3.5.4.26, 1.1.1.193, 3.1.3.104, 4.1.99.12, 2.5.1.78, 2.5.1.9, 2.7.1.26, 2.7.7.2). The isolate may be resistant to antibiotics due to the detection of CblA beta-lactamase (ARO:3002998). The 16S rRNA gene sequence of the species was most prevalent in the mouse gut (39.3% of 1,000 samples positive, at an average relative abundance of 1.51%), followed by wastewater (10.9%), and human gut (10.2%). The type strain is **NM69_E16B (=DSM110164**^**T**^). Its G+C content of genomic DNA is 46.0 mol%. It was isolated from the caecal/colon content of an APC^min/+^ Msh2^-/-^ mouse.

### Description of *Clostridium mucosae* sp. nov

*Clostridium mucosae* (mu.co’sae. N.L. gen. n. mucosae, of mucosa). The isolate shared the highest 16S rRNA gene sequence similarity with *Clostridium tertium* (99.22%), followed by *Clostridium sartagoforme* (98.17%). GTDB-Tk assigned the genome to a novel species within the genus *Clostridium*. This assignment was supported by the POCP value of 78.78% to *C. tertium* and 50.2% to *Clostridium butyricum*, the type species of this genus, and by the genome tree. The ANI and GGDC values between the genomes of the isolate and *C. tertium or C. sartagoforme* were below species delineation (91.01%/42.60% and 84.26%/28.40%, respectively), confirming that this isolate represents a novel species. The number of CAZymes identified within the genome was 307. Genome analysis predicted the ability to utilise glucose, arbutin, salicin, cellobiose, sucrose, trehalose, maltose, starch, and cellulose as carbon source. KEGG analysis identified pathways for the production of acetate from acetyl-CoA (EC:2.3.1.8, 2.7.2.1), propionate from propanoyl-CoA (EC:2.3.1.8, 2.7.2.1), L-cysteine and acetate from sulfide and L-serine (EC:2.3.1.30, 2.5.1.47), L-glutamate from ammonia via L-glutamine (EC:6.3.1.2, 1.4.1.-), folate (vitamin B9) from 7,8-dihydrofolate (EC:1.5.1.3), and riboflavin (vitamin B2) from GTP (EC:3.5.4.25, 3.5.4.26, 1.1.1.193, 3.1.3.104, 4.1.99.12, 2.5.1.78, 2.5.1.9, 2.7.1.26, 2.7.7.2). No antibiotic resistance genes were detected. The 16S rRNA gene sequence of the species was most prevalent in pig gut microbiota (60.4% of 1,000 samples positive), followed by activated sludge (52.6 %), and human gut microbiota (48.5%). The type strain is **PG-426-IM-1**^**T**^ **(=DSM100503**^**T**^). Its G+C content of genomic DNA is 27.7 mol%. It was isolated from the ileal mucosa of a TNF^deltaARE/+^ mouse.^12^

### Description of *Caniella* gen. nov

*Caniella* (Ca.ni.el’la. N.L. fem. n. *Caniella*, in honour of Prof. Dr. Patrice Cani, UCLouvain, Brussels, Berglium, for his contribution to the field of microbe-host interactions in metabolic diseases). The closest relatives based on 16S rRNA gene sequence identity are species within genus *Olsenella* (max. 94.36% to *Olsenella phocaeensis*). GTDB-Tk assigned the genomes to an unknown genus within family *Atopobiaceae*. The highest POCP value was to *Olsenella scatoligenes* (50.0-52.4%), whilst the values to *Olsenella uli* (type species of *Olsenella*) and to *Atopobium minutum* (type species of *Atopobium*) were 48.6-52.1% and 45.3-48.4%, respectively. Phylogenomic analysis confirmed that the isolates fall within the family *Atopobiaceae*, separated from members of the genera *Olsenella* and *Atopobium*. Together, these analyses support the novel genus classification of the isolates. The type species is *Caniella muris*.

### Description of *Caniella muris* sp. nov

*Caniella muris* (mu’ris L. gen. n. *muris* of a mouse). The species has all features of the genus. Cells usually grow as coccobacilli to short rods with slightly pointy ends (0.8-2.0 µm in length) in WCA medium under anaerobic conditions for 2-3 days. In total, 99-118 CAZymes were identified within the genome of strains within this species. Genome analysis predicted the ability to utilise arbutin, salicin, cellobiose, maltose, and starch. KEGG analysis identified the pathways for production of acetate from acetyl-CoA (EC:2.3.1.8, 2.7.2.1), propionate from propanoyl-CoA (EC:2.3.1.8, 2.7.2.1), L-cysteine and acetate from sulfide and L-serine (EC:2.3.1.30, 2.5.1.47), L-glutamate from ammonia via L-glutamine (EC:6.3.1.2, 1.4.1.-), and folate from 7,8-dihydrofolate (EC:1.5.1.3). No antibiotic resistance genes were detected. Ecological analysis suggests that the species is most prevalent within amplicon datasets from the mouse gut (ca. 17% of 1,000 samples positive). The range of G+C content of genomic DNA of strains within the species is 69.0-70.0 mol%. The type strain, **CLA-SR-94**^**T**^ **(=DSM 110323**^**T**^), was isolated from the caecal content of a wild mouse. Strain NM08_P-01 (=DSM 110563) was isolated from the caecal/colon content of an APC^min/+^ Msh2^-/-^ mouse. Strain CLA-SR-156 (=DSM 110983) and WCA-FA-Sto1.30.01 (=DSM 105314) were isolated in Aachen, Germany, from the caecal content of a conventional laboratory mouse and stomach content of a wild mouse, respectively.

### Description of *Dubosiella muris* sp. nov

*Dubosiella muris* (mu’ris L. gen. n. *muris* of a mouse). The closest relative to this isolate based on 16S rRNA gene sequence similarity is *Dubosiella newyorkensis* (91.10%), the type species of this genus. Despite this relatively low value that may indicate a novel genus, GTDB-Tk assigned the genome as a novel species within the genus *Dubosiella*. Both POCP and genome tree analysis further supported this genus classification, with a placement next to the aforementioned species and a corresponding POCP value of 73.82%, well above the genus delineation cut-off point of 50%. The ANI (77.66%) and GGDC (18.40%) values to the genome of *D. newyorkensis* confirmed that the isolate represents a novel species. The isolate grows in Phenylethyl Alcohol Medium under anaerobic conditions within 3 days. In total, 168 CAZymes were identified within the genome. The ability to utilise cellobiose and starch as carbon sources was predicted. KEGG analysis identified pathways for acetate production from acetyl-CoA (EC:2.3.1.8, 2.7.2.1), propionate from propanoyl-CoA (EC:2.3.1.8, 2.7.2.1), L-cysteine and acetate from sulfide and L-serine (EC:2.3.1.30, 2.5.1.47), and L-glutamate from ammonia via L-glutamine (EC:6.3.1.2, 1.4.1.-). The presence of the genes for tetracycline-resistant ribosomal protection protein (ARO:0000002) may indicate antibiotic resistance. The 16S rRNA gene sequence of the species was most prevalent in the mouse gut (14.8% of 1,000 samples positive, at an average relative abundance of 2.62%), followed by pig gut (2.5%), and human skin (1.7%). The type strain is **NM09_H32**^**T**^ **(=DSM 110160**^**T**^). Its G+C content of genomic DNA is 50.6 mol%. It was isolated from the caecal/colon content of an APC^min/+^ Msh2^-/-^ mouse.

### Description of *Flintibacter muris* sp. nov

*Flintibacter muris* (mu’ris L. gen. n. *muris* of a mouse). This bacterium shares highest 16S rRNA gene sequence identity to *Flintibacter butyricus* (97.32%), the type species of the genus *Flintibacter*. GTDB-Tk identified the genome as an unknown species within the genus *Lawsonibacter*. However, *Flintibacter* spp. are classified under the genus *Lawsonibacter* in GTDB, although this genus was validly published later.^85^ The highest POCP value (69.42%) is to *F. butyricus* and only 46.58% to *Lawsonibacter asaccharolyticus* (the type species of this genus). The genome tree analysis also identified *F. butyricus* as the closest relative. However, the ANI and GGDC value of 84.8% and 30.1%, respectively, between the latter species and the isolate confirmed its status of a novel species. The type strain forms rods when grown on BHI Medium under anaerobic conditions for 2 days. The total number of CAZymes identified in the genome was 165. The ability to utilise glucose and starch as carbon source was predicted. KEGG analysis identified pathways for acetate production from acetyl-CoA (EC:2.3.1.8, 2.7.2.1), propionate from propanoyl-CoA (EC:2.3.1.8, 2.7.2.1), L-cysteine and acetate from sulfide and L-serine (EC:2.3.1.30, 2.5.1.47), L-glutamate from ammonia via L-glutamine (EC:6.3.1.2, 1.4.1.-), and folate (vitamin B9) from 7,8-dihydrofolate (EC:1.5.1.3). As butyrate biosynthesis was not predicted, manual examination of the Prokka annotation identified genes assigned as 3-hydroxybutyryl-CoA dehydrogenase (EC 1.1.1.157), butyryl-CoA:acetate CoA-transferase (EC 2.8.3.-), 3-aminobutyryl-CoA aminotransferase (EC 2.6.1.111), and putative butyrate kinase 2 (EC 2.7.2.7). The presence of genes for tetracycline-resistant ribosomal protection protein (ARO:0000002) indicates antibiotic resistance. The 16S rRNA gene sequence of the species was most prevalent in the mouse gut (74.7% of 1,000 samples positive, at an average relative abundance of 0.98%), followed by chicken gut (71.5%), and pig gut (52.9%). The type strain is **CLA-AV-17**^**T**^ **(=DSM 110149** ^**T**^). Its G+C content of genomic DNA is 55.8 mol%. It was isolated from the caecal content of an SPF mouse.

### Description of *Hominisplanchenecus murintestinalis* sp. nov

*Hominisplanchenecus murintestinalis* (mur.in.tes.ti.na’lis. L. mas. or fem. n. *mus*, a mouse; N.L. masc. adj. *intestinalis*, intestinal; N.L. masc. adj. *murintestinalis*, of the mouse intestine). Strains CLA-AA-M05 and NM72_1-8 show highest 16S rRNA gene sequence identities to *Murimonas intestini* (91.57-91.13%), followed by *Marvinbryantia formatexigens* (91.39-91.47%). Phylogenomic analysis placed the genome of the isolates amongst members of multiple genera within family *Lachnospiraceae*. GTDB-Tk assigned the isolates to the genus ‘CAG-56’ (family *Lachnospiraceae*), the same genus as the recently described species *Hominisplanchenecus faecis*.^86^ The highest POCP value was 50.6-51.9% to *H. faecis* (DSM 113194), whilst that to *M. intestini* (type species) and *M. formatexigens* (type species) were 38.7-40.6% and 42.1-42.5%, respectively. However, ANI and GGDC values of the isolates to *H. faecis* were 71.4% and 20.5-22.9%, respectively, supporting the creation of a novel species to accommodate the isolates. Cells are rods (>2.0 µm in length), some are string like (longer than 10 µm), when grown in BHI medium under anaerobic conditions for 1-3 days. In total, 180-191 CAZymes were identified within the genomes. Gene prediction suggested the ability to utilise glucose and starch as carbon source. KEGG analysis identified pathways for the production of acetate from acetyl-CoA (EC:2.3.1.8, 2.7.2.1), propionate from propanoyl-CoA (EC:2.3.1.8, 2.7.2.1), L-glutamate from ammonia via L-glutamine (EC:6.3.1.2, 1.4.1.-), cobalamin (vitamin B12) from cobinamide (EC:2.5.1.17, 6.3.5.10, 6.2.1.10, 2.7.1.156), folate from 7,8-dihydrofolate (EC:1.5.1.3), and riboflavin (vitamin B2) from GTP (EC:3.5.4.25, 3.5.4.26, 1.1.1.193, 3.1.3.104, 4.1.99.12, 2.5.1.78, 2.5.1.9, 2.7.1.26, 2.7.7.2). No antibiotic resistance genes were detected. Ecological analysis suggested that the isolates are most prevalent within amplicon datasets from the mouse gut (41.6-42.2%) at an average relative abundance of 0.24%. The G+C content of genomic DNA is 45.8-45.9%. The type strain, **CLA-AA-M05**^**T**^ **(=DSM 111139**^**T**^), was isolated from the caecal content of a SPF mouse in Aachen, Germany. Strain NM72_1-8 (=DSM 110165) was isolated from an APC^min/+ Msh2-/-^ mouse in Toronto, Canada.

### Description of *Lactobacillus agrestimuris* sp. nov

*Lactobacillus agrestimuris* (a.gres.ti.mu’ris. L. masc. adj. *agrestis*, wild; L. masc. or fem. n. *mus*, a mouse; N.L. gen. n. *agrestimuris*, of a wild mouse). The isolate has the highest 16S rRNA gene sequence similarity to *Lactobacillus* species (max. 96.83% to *Lactobacillus hamsteri*). GTDB-Tk assigned the genome to an unknown species within the genus *Lactobacillus*. The POCP value to *L. hamsteri* (75.0%) and to *Lactobacillus delbrueckii* (60.2%), the type species of the genus *Lactobacillus*, further supports this genus placement. Genome tree analysis identified *L. hamsteri* as the closest relative. As none of the closely related *Lactobacillus* species with a valid name has an ANI value >95 % to the genome of the isolate, including *L. hamsteri* (ANI, 81.4%; GGDC, 21.6%), the creation of a novel species is required to accommodate the isolate. Cells were generally short rods (1.0 -2.0 µm) when grown on WCA medium under aerobic or anaerobic conditions. The total number of CAZyme in the genome was 124. Genome analysis predicted the ability to utilise glucose as carbon source. KEGG analysis identified pathways for acetate production from acetyl-CoA (EC:2.3.1.8, 2.7.2.1) and propionate from propanoyl-CoA (EC:2.3.1.8, 2.7.2.1). No antibiotic resistance genes were detected. The 16S rRNA gene sequence of the species was most prevalent in the chicken gut (84.8% of 1,000 samples positive, at an average relative abundance of 9.94%), followed by human vagina (84.4%), and pig gut (68.1%). The type strain is **CLA-SR-99** ^**T**^ **(=DSM110155**^**T**^). Its G+C content of genomic DNA is 34.8 mol%. It was isolated from the caecal content of a wild mouse.

### Description of *Lactococcus muris* sp. nov

*Lactococcus muris* (mu’ris L. gen. n. *muris* of a mouse). Based on 16S rRNA gene sequence analysis (883 bp), the isolate is phylogenetically related to multiple species within the genus *Lactococcus*, with highest sequence identity to *Lactococcus garvieae* subsp. *garvieae* (96.72%). GTDB-Tk classified the genome as “Lactococcus sp002492185”. The highest POCP value was 73.72 % to *L. garvieae* and that to the type species of the genus, *L. lactis*, was 61.4%. This together with genome tree analysis confirmed classification of the isolate within the genus *Lactococcus*. The status of a novel species was confirmed by ANI (82.6%) and GGDC (26.7-26.8%) values to *L. garvieae*. The total number of CAZymes identified in the genome was 142. Genome analysis predicted the ability to utilise arbutin, salicin, cellobiose, and starch as carbon sources. KEGG analysis identified pathways for acetate production from acetyl-CoA (EC:2.3.1.8, 2.7.2.1), propionate from propanoyl-CoA (EC:2.3.1.8, 2.7.2.1), L-cysteine and acetate from sulfide and L-serine (EC:2.3.1.30, 2.5.1.47), and folate (vitamin B9) from 7,8-dihydrofolate (EC:1.5.1.3) No antibiotic resistance genes were identified. The 16S rRNA gene sequence of the species was most prevalent in wastewater (11.1% of 1,000 samples positive), followed by pig gut (8.8%), and activated sludge (6.5%). The type strain is **HZI-1**^**T**^ **(=DSM 109779**^**T**^). Its G+C content of genomic DNA is 39.2 mol%. It was isolated from the caecal content of an SPF mouse.

### Description of *Lactococcus ileimucosae* sp. nov

*Lactococcus ileimucosae (i*.*le*.*i*.*mu*.*co’sae. L. neut. n. ileum*, ileum; *N*.*L. fem*.*n. mucosa*, mucosa; *N*.*L. gen. n. ileimucosae*, of ileal mucosa, the mouse tissue used for isolation). The closest relatives based on 16S rRNA gene sequences were species within the genus *Lactococcus* (max. 98.14% to *Lactococcus formosensis*). GTDB-Tk classified the genome as an unknown species within the genus *Lactococcus*. The highest POCP value was to *L. formosensis* (74.8%), and that to *L. lactis* (the type species of this genus) was 61.0%, supporting assignment within the genus *Lactococcus*. This was confirmed by genome tree analysis. As none of the closest relatives shared ANI and GGDC values above the corresponding species delineation thresholds of 95% and 70%, respectively, including *L. formosensis* (ANI, 82.06%; GGDC, 26.50%) and *Lactococcus garvieae* subsp. *garvieae* (a close relative in the genome tree; ANI, 82.16%; GGDC, 32.10%), a novel species is proposed to accommodate this isolate. Cells grow as spindle-shaped coccobacilli (ca. 1.0 µm in length) in BHI medium under anaerobic conditions. The number of CAZymes identified in the genome was 146. Genome analysis predicted the ability to utilise arbutin, salicin, cellobiose, starch as carbon source. KEGG analysis identified pathways for the production of acetate from acetyl-CoA (EC:2.3.1.8, 2.7.2.1), propionate from propanoyl-CoA (EC:2.3.1.8, 2.7.2.1), L-cysteine and acetate from sulfide and L-serine (EC:2.3.1.30, 2.5.1.47), and folate (vitamin B9) from 7,8-dihydrofolate (EC:1.5.1.3). No antibiotic resistance genes were detected. The 16S rRNA gene sequence of the species was most prevalent in the pig gut (8.9% of 1,000 samples positive), followed by wastewater (7.7%), and insect gut (6.2%). The type strain is **M9-GB-M-SO-A**^**T**^ **(=DSM 107391**^**T**^). Its G+C content of genomic DNA is 39.5 mol%. It was isolated from the jejunal/ileal mucosa of a wild mouse.

### Description of *Lepagella* gen. nov

*Lepagella* (Le.pa.gel’la. N.L. fem. n. *Lepagella*, in honour of Dr. Patricia Lepage, INRAE, Jouy-en-Josas, France, for her contribution to the field of gut microbiome research in health and disease). The closest neighbours based on 16S rRNA gene sequence comparison are species within family *Muribaculaceae* (max. 86.91% to *Duncaniella freteri*). GTDB-Tk assigned the isolate to the species ‘sp002493045’, within the yet unknown genus ‘CAG-485’ (family *Muribaculaceae*). Phylogenetic analysis identified the genome to be in the same clade as *Muribaculum intestinale*. However, none of the close relatives with a valid name was characterized by a POCP value >50 %, including *M. intestinale* (type genus of the family; 41.9%) and *Duncaniella muris* (44.4%). This data supports the creation of a novel genus to accommodate the isolate. The type species is *Lepagella muris*.

### Description of *Lepagella muris* sp. nov

*Lepagella muris* (mu’ris L. gen. n. *muris* of a mouse). The species has all features of the genus. It grows in Columbia Blood Medium (5 % sheep blood) under anaerobic conditions within 3 days. In total, 371 CAZYmes were identified within the genome. Gene prediction revealed the ability to utilise starch as carbon source. KEGG analysis identified pathways for the production of acetate from acetyl-CoA (EC:2.3.1.8, 2.7.2.1), propionate from propanoyl-CoA (EC:2.3.1.8, 2.7.2.1), L-cysteine and acetate from sulfide and L-serine (EC:2.3.1.30, 2.5.1.47), L-glutamate from ammonia via L-glutamine (EC:6.3.1.2, 1.4.1.-), cobalamin (vitamin B12) from cobinamide (EC:2.5.1.17, 6.3.5.10, 6.2.1.10, 2.7.1.156), and folate (vitamin B9) from 7,8-dihydrofolate (EC:1.5.1.3). Antibiotic resistance may be conferred via tetracycline-resistant ribosomal protection protein (ARO:0000002). The 16S rRNA gene sequence of the species was most prevalent in the mouse gut (31.1% of 1,000 samples positive, mean rel. abund. 2.73%), followed by human skin (5.1%), and pig gut (3.0%). The type strain is **NM04_E33**^**T**^ **(=DSM 110157**^**T**^). Its G+C content of genomic DNA is 46.1 mol%. It was isolated from caecal/colon content of an APC^min/+^ Msh2^-/-^ mouse.

### Description of *Limosilactobacillus caecicola* sp. nov

*Limosilactobacillus caecicola* (cae.ci’co.la. L. neut. n. *caecum*, caecum;L. masc./fem. suff. *-cola*, dweller; from L. masc./fem. n. *incola*, inhabitant, dweller; N.L. n. *caecicola*, an inhabitant of the caecum). The closest relative based on 16S rRNA gene sequence identity is *Limosilactobacillus coleohominis* (98.66%). GTDB-Tk classified the genome as an unknown species within the genus *Limosilactobacillus*. The highest POCP value was 76.0 % to *L. coleohominis*, whilst that to *L. fermentum* (the type species of the genus *Limosilactobacillus*) was 66.9%. The genome tree also placed the bacterium within limosilactobacilli next to *L. coleohominis*. Genome comparison to the type strain of the latter species using ANI and GGDC indicates that the isolate represents a novel species (79.9% and 19.3%, respectively). Cells are straight to slightly curved rods (1-2 µm in length) when grown on WCA medium under anaerobic conditions for 2-3 days. In total, 115 CAZymes were identified within the genome. KEGG analysis identified pathways for acetate production from acetyl-CoA (EC:2.3.1.8, 2.7.2.1), and propionate from propanoyl-CoA (EC:2.3.1.8, 2.7.2.1). No carbon source utilisation (of those tested in Protologger) and antibiotic resistance genes were predicted. The 16S rRNA gene sequence of the species was most prevalent in the chicken gut (32.5% of 1,000 samples positive), followed by pig gut (27.8%), and human vagina (9.9%). The type strain is **CLA-SR-145**^**T**^ **(=DSM 110982**^**T**^). Its G+C content of genomic DNA is 44.7 mol%. It was isolated from the caecal content of a wild mouse.

### Description of *Limosilactobacillus agrestimuris* sp. nov

*Limosilactobacillus agrestimuris* (a.gres.ti.mu’ris. L. masc. adj. *agrestis*, wild; L. masc. or fem. n. *mus*, a mouse; N.L. gen. n. agrestimuris, of a wild mouse). The isolate shares highest 16S rRNA gene sequence identities to species within the genus *Limosilactobacillus* (max. 99.87% to *Limosilactobacillus urinaemulieris*). GTDB-Tk classified the genome as an unknown species within the genus *Limosilactobacillus*. The highest POCP value was to *Limosilactobacillus vaginalis* (82.6%), whilst value to *Limosilactobacillus fermentum* (the type species of this genus) was 63.3%. This supports placement of the isolate within the genus *Limosilactobacillus*. Genome tree analysis placed the isolate next to *L. vaginalis*. However, none of the close relatives (including *L. urinaemulieris* and *L. vaginalis*) share ANI and GGDC values above the species delineation cut-off points, confirming the status of this isolate as a novel species within the genus *Limosilactobacillus*. The bacterium grows well on WCA medium under anaerobic conditions within 1-3 days. The total number of CAZymes identified in the genome was 118. Genome analysis could not identify any genes related to carbon source utilisation, but predicted the ability to produce acetate, propionate, and folate. No antibiotic genes were detected.

The 16S rRNA gene sequence of the species was most prevalent in the chicken gut (91.2% of 1,000 samples positive, at an average relative abundance of 2.63%), followed by pig gut (81.0%), and mouse gut (40.5%). The type strain is **WCA-sto-4**^**T**^ **(=DSM 106037**^**T**^). Its G+C content of genomic DNA is 39.7 mol%. It was isolated from the gut content of a wild mouse.

### Description of *Mediterraneibacter agrestimuris* sp. nov

*Mediterraneibacter agrestimuris* (a.gres.ti.mu’ris. L. masc. adj. *agrestis*, wild; L. masc. or fem. n. *mus*, a mouse; N.L. gen. n. *agrestimuris*, of a wild mouse). The closest phylogenetic neighbours to the isolate based on 16S rRNA gene sequences was *Ruminococcus torques* (96.74%), followed by *Faecalicatena contorta* (96.31%) and *Mediterraneibacter glycyrrhizinilyticus* (96.31%). GTDB-Tk assigned the genome as an unknown species under the genus *Muricomes*. However, the POCP value to *Muricomes intestini* (type species), *Faecalicatena contorta* (type species), *Ruminococcus torques*, and *Ruminococcus flavefaciens* (type species) were 36.5%, 45.8%, 47.8%, and 22.7%, respectively, all below the genus delineation threshold. The highest POCP value was to *M. glycyrrhizinilyticus* (56.23%), and the value to *Mediterraneibacter massiliensis* (the type species of the genus *Mediterraneibacter*) was 50.11%. The genome tree analysis also identified *M. glycyrrhizinilyticus* as the closest relative. Moreover, the 16S rRNA gene sequence identity between the isolate and *M. intestini* is only 93.93%. These analyses indicate that the isolate belongs to the genus *Mediterraneibacter* and not *Muricomes*. None of the closely related species (including *M. glycyrrhizinilyticus*, ANI: 78.9%, GGDC: 21.5%) had ANI and GGCD values above the species cut-off value, confirming the status of this isolate as a novel species. Cells grow as rods (ca. 1.8-3.0 µm in length) in WCA medium under anaerobic conditions for 1-3 days. The total number of CAZymes identified in the genome was 263. Genome analysis predicted the ability to utilise starch, cellulose, sulfide, and L-serine. The genes for production of acetate, propionate, L-cysteine, L-glutamate, and folate were detected. The presence of the genes for tetracycline-resistant ribosomal protection protein (ARO:0000002), *vanR* (ARO:3000574), and *vanS* (ARO:3000071) suggests antibiotic resistance. The 16S rRNA gene sequence of the species was most prevalent in wastewater (34.4% of 1,000 samples positive), followed by the human gut (27.5%); the prevalence in mouse gut was 11.0% (average relative abundance, 0.34%). The type strain is **CLA-SR-176**^**T**^ **(=DSM 111629**^**T**^). Its G+C content of genomic DNA is 41.9 mol%. It was isolated from the caecal content of a wild mouse.

### Description of *Muribaculum caecicola* sp. nov

*Muribaculum caecicola* (cae.ci’co.la. L. neut. n. *caecum*, caecum; L. masc./fem. suff. *-cola*, dweller; from L. masc./fem. n. *incola*, inhabitant, dweller; N.L. n. *caecicola*, an inhabitant of the caecum). The isolate shares closest 16S rRNA gene identity to *Muribaculum intestinale* (90.77%), the type species of the genus *Muribaculum*. GTDB-Tk identified the genome as ‘Muribaculum sp002473395’ under the genus *Muribaculum*. Both the POCP value of 57.0% to *M. intestinale* and topology of the genomic tree support this genus-level classification. The ANI and GGDC value to *M. intestinale* were 71.1% and 39.3%, respectively, confirming the novel species status of this isolate. It grows in Anaerobic Brain Heart Infusion under anaerobic conditions within 24 h. The total number of CAZymes identified in the genome was 189. Genome analysis predicted the ability to utilise starch and to produce acetate, propionate, L-glutamate, and riboflavin (vitamin B2). No antibiotic resistance genes were identified. The 16S rRNA gene sequence of the species was most prevalent in the mouse gut (27.4% of 1,000 samples positive). The type strain is **NM86_A22**^**T**^ **(=DSM 110169**^**T**^). Its G+C content of genomic DNA is 45.7 mol%. It was isolated from the caecal/colon content of an APC^min/+^ Msh2^-/-^ mouse.

### Description of *Muricaecibacterium* gen. nov

*Muricaecibacterium* (Mu.ri.cae.ci.bac.te.ri.um. L. masc. n. mus, a mouse; L. neut. n. caecum, the caecum; bacterium, L. neut. n. a bacterium; Muricaecibacterium, a microbe from the caecum of a mouse). The closest phylogenetic neighbours based on 16S rRNA gene sequence similarity are species within the genus *Olsenella* (max. 92.78% to *Olsenella umbonata*). GTDB-Tk classified the isolate as an unknown genus within the family *Atopobiaceae*. The highest POCP value was 52.1% to *Olsenella uli*, the type species of the genus *Olsenella*, whilst 48.2 % to *Atopobium minutum*, the type species of the genus *Atopobium*. Although the genome tree analysis placed the isolate within the family *Atopobiaceae*, it branched separately from members of the genera *Olsenella* and *Atopobium*. Together with the GTDB-Tk assignment aforementioned, this supports the creation of a novel genus status for the isolate. The type species is *Muricaecibacterium torontonense*.

### Description of *Muricaecibacterium torontonense* sp. nov

*Muricaecibacterium torontonense* (N.L. neut. adj. *torontonense*, pertaining to Toronto (Canada), where the bacterium was isolated). The species has all features of the genus. Cells usually grow singly, in pairs or in short serpentine chains in Sulfite Polymyxin Sulfadiazine medium under anaerobic conditions for up to 3 days. In total, 94 CAZymes were identified within the genome of the type strain. Further genome analyses predicted the ability to utilise arbutin, salicin, cellobiose, maltose, and starch. KEGG analysis identified the pathways for production of acetate from acetyl-CoA (EC:2.3.1.8, 2.7.2.1), propionate from propanoyl-CoA (EC:2.3.1.8, 2.7.2.1), L-glutamate from ammonia via L-glutamine (EC:6.3.1.2, 1.4.1.-), and folate (vitamin B9) from 7,8-dihydrofolate (EC:1.5.1.3). No antibiotic resistance genes were detected. The 16S rRNA gene sequence of the species was most prevalent in the mouse gut (10.5% of 1,000 samples positive). The type strain is **NM07_P-09**^**T**^ **(=DSM 110159**^**T**^). Its molecular G+C content of genomic DNA is 58.8 mol%. It was isolated from the caecal content of an APC^min/+^ Msh^2-/-^ mouse.

### Description of *Neobacillus muris* sp. nov

*Neobacillus muris* (mu’ris L. gen. n. *muris* of a mouse). The closest phylogenetic neighbour to the isolate based on 16S rRNA gene sequence similarity is *Neobacillus drentensis* (98.22%). GTDB-Tk assignment to the genus *Neobacillus* was confirmed by POCP values >50% to multiple *Neobacillus* spp., including *N. drentensis* (64.37%) and *Neobacillus cucumis* (65.54%). In addition to a lack of species assignment by GTDB-Tk, comparison to all close relatives provided ANI values below 95%, supporting the assignment of a novel species within the genus *Neobacillus*. The isolate was observed to have a large CAZyme repertoire, containing 348 CAZymes along with the pathways for utilisation of glucose, trehalose, maltose, and starch. The antibiotic resistance genes, vanR (ARO:3000574), vanZ (ARO:3000116), and vanS (ARO:3000071) were detected within the genome, along with a major facilitator superfamily (MFS) antibiotic efflux pump (ARO:0010002). Acetate, butyrate, and propionate were all predicted to be produced. Genome analysis also identified the presence of pathways for the production of cobalamin (vitamin B12) from cobinamide (EC:2.5.1.17, 6.3.5.10, 6.2.1.10, 2.7.1.156), folate (vitamin B9) from 7,8-dihydrofolate (EC:1.5.1.3) and riboflavin (vitamin B2) from GTP (EC:3.5.4.25, 3.5.4.26, 1.1.1.193, 3.1.3.104, 4.1.99.12, 2.5.1.78, 2.5.1.9, 2.7.1.26, 2.7.7.2). The 16S rRNA gene sequence of the species was most prevalent in the rhizosphere (71.2% of 1,000 samples positive), followed by soil (55.2%), and plant metagenomic samples (49.1%). The type strain is **CLA-SR-152**^**T**^ **(=DSM 110989**^**T**^). Its G+C content of genomic DNA is 41.6 mol%. It was isolated from caecal content of an SPF mouse.

### Description of *Odoribacter lunatus* sp. nov

*Odoribacter lunatus* (lu.na’tus. L. masc. adj. *lunatus*, crescent-shaped, pertaining to the cell shape). According to 16S rRNA gene sequence analysis, the isolate is most closely related to *Odoribacter laneus* (90.05%). GTDB-Tk assigned the genome as an unknown species within the genus *Odoribacter*. The highest POCP value was to *Odoribacter laneus* (55.2%), whereas the value to *Odoribacter splanchnicus* (the type species of this genus) was 47.5%. The genome tree also placed the isolate within a monophyletic clade with other two *Odoribacter* species. The ANI and GGDC value of the isolate to these two *Odoribacter* species were all well below the species delineation cut-off point, therefore confirming the novel status of the isolate within the genus *Odoribacter*. Cells are rods with pointy ends forming a crescent shape (length: ca. 0.8-1.5 µm) when grown on mGAM agar under anaerobic conditions for 3-10 days. The total number of CAZymes identified in the genome was 187. Genome analysis could not find any genes related to carbon source utilisation, but identified the genes for production of acetate (from acetyl-CoA, EC:2.3.1.8, 2.7.2.1), propionate (from propanoyl-CoA, EC:2.3.1.8, 2.7.2.1), L-glutamate (via L-glutamine, EC:6.3.1.2, 1.4.1.-), and riboflavin (vitamin B2; from GTP, EC:3.5.4.25, 3.5.4.26, 1.1.1.193, 3.1.3.104, 4.1.99.12, 2.5.1.78, 2.5.1.9, 2.7.1.26, 2.7.7.2). No antibiotic resistance genes were identified. The ecological analysis based on 16S rRNA gene amplicons could not identify any ecosystem with the presence of this species. The type strain is **CLA-AA-M09**^**T**^ **(=DSM 112344**^**T**^). Its G+C content of genomic DNA is 43.2 mol%. It was isolated from a filtered (0.45 µm) caecal slurry from a wild mouse.

### Description of *Otoolea* gen. nov

*Otoolea* (O.too’le.a. N.L. fem. n. *Otoolea*, in honour of Prof. Dr. Paul O’Toole, University College Cork, Ireland, for his contribution to the field of gut microbiome research). The isolate showed highest 16S rRNA gene sequence similarities to species with family *Lachnospiraceae* (max. 94.20% to *Clostridium fessum*; classified under family *Clostridiaceae* in LPSN). GTDB-Tk assigned the genome to the genus ‘Clostridium_Q’ within family *Lachnospiraceae*. The phylogenetic analysis identified the strain to be in a clade containing *Clostridium* species. However, none of the closest relatives shared a POCP value above the genus delineation value (50 %), with a maximum value of 46.7% to *C. fessum*, followed by *C. symbiosum* (45.3%). POCP value to *Clostridium butyricum*, the type species of this genus, was only 20.8 %. This data supports the creation of a novel genus to accommodate the isolate. The type species is *Otoolea muris*.

### Description of *Otoolea muris* sp. Nov

*Otoolea muris* (mu’ris L. gen. n. *muris* of a mouse). The species has all features of the genus. Cells are rods (1.0-5.0 µm in length) when grown in BHI medium under anaerobic conditions for 24 hours. In total, 310 CAZymes were identified in the genome. Gene prediction revealed the ability to utilise starch as carbon source. KEGG analysis identified pathways for the production of acetate from acetyl-CoA (EC:2.3.1.8, 2.7.2.1), propionate from propanoyl-CoA (EC:2.3.1.8, 2.7.2.1), L-cysteine and acetate from sulfide and L-serine (EC:2.3.1.30, 2.5.1.47), L-glutamate from ammonia via L-glutamine (EC:6.3.1.2, 1.4.1.-), cobalamin (vitamin B12) from cobinamide (EC:2.5.1.17, 6.3.5.10, 6.2.1.10, 2.7.1.156), and riboflavin (vitamin B2) from GTP (EC:3.5.4.25, 3.5.4.26, 1.1.1.193, 3.1.3.104, 4.1.99.12, 2.5.1.78, 2.5.1.9, 2.7.1.26, 2.7.7.2). No antibiotic resistance genes were detected. The 16S rRNA gene sequence of the species was most prevalent in the mouse gut (54.5% of 1,000 samples positive), followed by chicken gut (17.3%), and pig gut (12.7%). The type strain is **CLA-AA-M04**^**T**^ **(=DSM 111138**^**T**^). Its G+C content of genomic DNA is 50.8 mol%. It was isolated from the caecal content of an SPF mouse.

### Description of *Palleniella muris* sp. nov

*Palleniella muris* (mu’ris L. gen. n. *muris* of a mouse). Based on previous analyses, 16S rRNA gene sequence similarities between members of the family *Prevotellaceae* have been shown to be uninformative for the placement of novel isolates; this data is thus not included here.^87^ Phylogenomic placement of the type strain identified it as a member of the genus *Palleniella*, placed next to the type species of this genus, *Palleniella intestinalis*. ANI analysis to all members of *Palleniella*, and the neighbouring genera (*Xylanibacter, Leyella, Hoylesella, Segatella, Hallella*, and *Prevotella*), confirmed that the type strain represents a novel species as all values were below the species delineation threshold of 95%. The highest value was to *P. intestinalis* (92.05%). The isolate grows in Anaerobic Brain Heart Infusion under anaerobic conditions within 3 days. In the genome, 306 CAZymes were identified along with pathways for the utilisation of starch. KEGG analysis identified pathways for the production of acetate from acetyl-CoA (EC:2.3.1.8, 2.7.2.1), propionate from propanoyl-CoA (EC:2.3.1.8, 2.7.2.1), L-glutamate from ammonia via L-glutamine (EC:6.3.1.2, 1.4.1.-), and folate (vitamin B9) from 7,8-dihydrofolate (EC:1.5.1.3). Ecological analysis suggests that the species is most prevalent within amplicon datasets from the mouse gut (2.3%) at an average relative abundance of 3.29%. The type strain is **NM73_A23**^**T**^ **(=DSM 110166**^**T**^). Its G+C content of genomic DNA is 47.1 mol%. It was isolated from the caecal/colon content of an APC^min/+^ Msh2^-/-^ mouse.

### Description of *Parasutterella muris* sp. nov

*Parasutterella muris* (mu’ris L. gen. n. *muris* of a mouse). The closest relative to strain CLA-SR-150, CLA-RA-1, and NM82_D38 based on 16S rRNA gene sequence identity was *Parasutterella excrementihominis* (95.75-96.38%). GTDB-Tk classified the genome as an unknown species within the genus *Parasutterella*. The highest POCP value was to *P. excrementihominis* (the type species of this genus; 74.1-78.8%), followed by *Turicimonas muris* (71.4-74.5%). The genome tree analysis also identified *P. excrementihominis* as the closest relative. However, the ANI and GGDC values to both *P. excrementihominis* and *T. muris* were all well below the species delineation cut-offs, justifying the proposal to create a novel species within the genus *Parasutterella* to accommodate these isolates. Cells are rods (1.0-1.2 µm in length) when grown in WCA medium under anaerobic conditions for 1-5 days. The total number of CAZymes identified in the genomes were 91-94. Genome analysis predicted the ability to utilise sulfide and L-serine for production of L-cysteine and acetate (EC:2.3.1.30, 2.5.1.47). No genes for carbon source utilisation were found. The 16S rRNA gene sequence of the species was most prevalent in the mouse gut (27.3-27.9% of 1,000 samples positive), followed by chicken gut (3.6-4.1%). The range of G+C content of genomic DNA of strains within this species is 48.9-49.4 mol%. The type strain is **CLA-SR-150**^**T**^ **(=DSM 111000**^**T**^). It was isolated from the caecal content of a conventionally colonized laboratory mouse. Strain CLA-RA-1 (=DSM 108034) and NM82_D38 (=DSM 110635) were isolated from the gut content of a wild mouse and the caecal/colon content of an APC^min/+^ Msh2^-/-^ mouse, respectively.

### Description of *Petralouisia* gen. nov

*Petralouisia* (Pe.tra.lou.i’si.a. N.L. fem. n. *Petralouisia*, in honour of Dr. Petra Louis, Rowett Institute, Abderdeen, Scotland, for her contribution to the field of gut microbiology). Based on 16S rRNA gene sequence similarities, the closest relatives to the isolate are *Ruminococcus gnavus* (90.93%), *Roseburia inulinivorans* (90.83%), and *Enterocloster aldensis* (90.80%). GTDB-Tk assigned the isolate to the unknown genus ‘g 14-2’ within family *Lachnospiraceae*. Phylogenomic analysis confirmed that this genus falls within family *Lachnospiraceae* between members of the genera *Eubacterium* and *Pseudobutyrivibrio*. All POCP values to the closest relatives were below 50%, supporting the creation of a novel genus to accommodate this isolate. The type species is *Petralouisia muris*.

### Description of *Petralouisia muris* sp. nov

*Petralousia muris* (mu’ris L. gen. n. *muris* of a mouse). The species shares all features of the genus. It grows in Anaerobic Brucella Medium supplemented with blood under anaerobic conditions within 4 days. In total, 316 CAZymes were identified within the genome of the type strain. Glucose, arbutin, salicin, trehalose, starch, and cellulose were predicted to be utilised carbon sources. KEGG analysis identified pathways for the production of acetate from acetyl-CoA (EC:2.3.1.8, 2.7.2.1), propionate from propanoyl-CoA (EC:2.3.1.8, 2.7.2.1), L-cysteine and acetate from sulfide and L-serine (EC:2.3.1.30, 2.5.1.47), and L-glutamate from ammonia via L-glutamine (EC:6.3.1.2, 1.4.1.-). Genome analysis also identified the presence of pathways for the production of cobalamin (vitamin B12) from cobinamide (EC:2.5.1.17, 6.3.5.10, 6.2.1.10, 2.7.1.156), folate (vitamin B9) from 7,8-dihydrofolate (EC:1.5.1.3), and riboflavin (vitamin B2) from GTP (EC:3.5.4.25, 3.5.4.26, 1.1.1.193, 3.1.3.104, 4.1.99.12, 2.5.1.78, 2.5.1.9, 2.7.1.26, 2.7.7.2). Antibiotic resistance may be conferred by expression of the glycopeptide resistance gene cluster vanR (ARO:3000574). Ecological analysis suggested that the species is most prevalent within amplicon datasets from the mouse gut (36.0%) at an average relative abundance of 0.85%. The type strain is **NM01_1-7b**^**T**^ **(=DSM 110156**^**T**^). Its G+C content of genomic DNA is 44.0 mol% It was isolated from the caecal/colon content of an APC^min/+^ Msh2^+/-^ mouse.

### Description of *Pumilibacteraceae* fam. nov

*Pumilibacteraceae* (Pu.mi.li.bac.te.ra.ce’ae. N.L. masc. n. *Pumilibacter*, type genus of the family; L. fem. pl. suff. *-aceae*, ending to denote a family; N.L. fem. pl. n. *Pumilibacteraceae*, the family of the genus Pumilibacter). The closest phylogenetic relatives based on 16S rRNA gene similarities are *Christensenella, Caldicoprobacter*, and *Saccharofermentans* spp. (<85.5%) within the order *Eubacteriales*. Phylogenomic analysis confirmed that the isolates form a monophyletic group distinct from all close relatives. The creation of a novel family was further supported by GTDB-Tk placement as ‘f_CAG-552’ within the order ‘*Christensenellales’* (not valid). Taxonomic classification of these bacteria at the order level, and the corresponding nomenclature, will require amendments in the future. Members of this new family were identified to be prevalent within the gastrointestinal tract of mice, although at sub-dominant levels (mean relative abundance <0.5%). The type genus of this family is *Pumilibacter*.

### Description of *Pumilibacter* gen. nov

*Pumilibacter* (Pu.mi.li.bac.ter. L. masc. n. *pumilus*, dwarf; N.L. masc. n. *bacter*, rod, referring to a bacterium in biology; N.L. masc. n. *Pumilibacter*, dwarf bacterium, pertaining to the small size of the type species). The closest phylogenetic relative based on 16S rRNA gene sequence identity are *Saccharofermentans acetigenes* (83.99%, to CLA-AA-M08) and *Christensenella hongkongensis* (85.65%, to CLA-AA-M10). POCP values to all close relatives were below 30% and GTDB-Tk placement assigned the type species to an unknown genus within family ‘f_CAG-552’. The type species is *Pumilibacter muris*.

### Description of *Pumilibacter intestinalis* sp. nov

*Pumilibacter intestinalis* (in.tes.ti.na’lis. N.L. fem adj. *intestinalis*, pertaining to the intestine). The species has all features of the genus. Additional phylogenetic relatives based on 16S rRNA gene sequences are *Ruminiclostridium josui* (84.32%), *Ruminiclostridium cellulolyticum* (84.11%), and *Vallitalea guaymasensis* (83.95%). Assignment to *the genus Pumilibacter* was confirmed by a POCP value of 69.7% between the genome of the type strain and that of the type species of the genus, *Pumilibacter muris*. Cells are rods (ca. 2.0 µm in length) to long rods (>5.0 µm in length) with a diameter of ca. 0.4 µm when grown on YCFA or mGAM blood agar under anaerobic conditions for 7 days. In total, 155 CAZymes were identified within the genome and only starch was predicted to be utilised as a carbon source. KEGG analysis identified pathways for the production of acetate from acetyl-CoA (EC:2.3.1.8, 2.7.2.1), propionate from propanoyl-CoA (EC:2.3.1.8, 2.7.2.1), L-cysteine and acetate from sulfide and L-serine (EC:2.3.1.30, 2.5.1.47), and L-glutamate from ammonia via L-glutamine (EC:6.3.1.2, 1.4.1.-). Antibiotic resistance was predicted based on the detection of tetracycline-resistant ribosomal protection protein. Ecological analysis suggested that the species is most prevalent within amplicon datasets from the mouse gut (39.9%). The type strain is **CLA-AA-M10**^**T**^. Its G+C content of genomic DNA is 49.6%. It was isolated from a filtered (0.45 µm) caecal suspension of an SPF mouse.

### Description of *Pumilibacter muris* sp. nov. 1

*Pumilibacter muris* (mu’ris L. gen. n. *muris* of a mouse). The species has all features of the genus. Additional phylogenetic relatives based on 16S rRNA gene sequences are X*ylanivirga thermophila* (83.77%), *Ruminiclostridium josui* (83.68%), and *Hespellia porcina* (83.57%). Separation from the other novel species within this genus represented by strain CLA-AA-M10 (described below) was confirmed via an ANI value of 76.4% and GGDC value of 25% between the two genomes. Cells are very small and spherical (diameter: 0.3-0.5 µm) when grown on YCFA agar under anaerobic conditions for 7 days. In total, 121 CAZymes were identified within the genome of the type strain and only starch was predicted to be utilised as a carbon source. KEGG analysis identified pathways for the production of acetate from acetyl-CoA (EC:2.3.1.8, 2.7.2.1), propionate from propanoyl-CoA (EC:2.3.1.8, 2.7.2.1), L-cysteine and acetate from sulfide and L-serine (EC:2.3.1.30, 2.5.1.47), and L-glutamate from ammonia via L-glutamine (EC:6.3.1.2, 1.4.1.-). No antibiotic resistance genes were identified within the genome. Ecological analysis suggested that the species is most prevalent within amplicon datasets from the mouse gut (32.8%). The type strain is **CLA-AA-M08**^**T**^. Its G+C content of genomic DNA is 46.81%. It was isolated from a filtered (0.45 µm) caecal suspension of an SPF mouse.

### Description of *Senimuribacter* gen. nov

*Senimuribacter* (Se.ni.mu.ri.bac.ter. L. masc. adj. *senex*, old; L. masc. n. or fem. *mus*, a mouse; N.L. masc. n. *bacter*, rod; N.L. masc. n. *Senimuribacter*, rod-shaped bacterium isolated from old mouse). The closest relatives based on 16S rRNA gene similarity are members of the genus *Eubacterium* (*Eubacterium sulci*, 91.54-92.20%, *Eubacterium infirmum*, 91.30-92.20%) and *Aminipila* (*Aminipila butyrica*, 90.74-91.77%). Phylogenomic analysis indicated that the isolate falls between members of the genera *Eubacterium* and *Mogibacterium*. The creation of a novel genus was further supported by GTDB-Tk placement as ‘g_Emergencia’, a genus proposed in 2016 but never validated.^88^ POCP values to all close relatives were <40%, greatly below the genus delineation threshold. The type species is *Senimuribacter intestinalis*.

### Description of *Senimuribacter intestinalis* sp. nov

*Senimuribacter intestinalis* (in.tes.ti.na’lis. N.L. fem adj. *intestinalis*, pertaining to the intestine). The species shares all features of the genus. The description of this species is based on two strains, YCFAG-7-CC-SB-Schm-I and C1.7. Cells are rods (0.8-1.7 µm in length) when grown in WCA medium under anaerobic conditions for 2-5 days. Strains of this species contain 102-113 CAZymes within their genome but no carbohydrate utilisation pathways were identified. KEGG analysis identified pathways for the production of acetate from acetyl-CoA (EC:2.3.1.8, 2.7.2.1), butyrate from butanoyl-CoA (EC:2.8.3.8), propionate from propanoyl-CoA (EC:2.3.1.8, 2.7.2.1), L-cysteine and acetate from sulfide and L-serine (EC:2.3.1.30, 2.5.1.47), folate (vitamin B9) from 7,8-dihydrofolate (EC:1.5.1.3), and riboflavin (vitamin B2) from GTP (EC:3.5.4.25, 3.5.4.26, 1.1.1.193, 3.1.3.104, 4.1.99.12, 2.5.1.78, 2.5.1.9, 2.7.1.26, 2.7.7.2). Ecological analysis suggested that the species is most prevalent within amplicon datasets from the mouse gut (ca. 51%). The type strain is **YCFAG-7-CC-SB-Schm-I**^**T**^ **(=DSM 106208**^**T**^). Its G+C content of genomic DNA is 43.9 mol%. It was isolated from caecal content of a 40-week-old SPF mouse in Freising, Germany. Strain C1.7 (=DSM 109599) was isolated from caecal content of an SPF mouse in Braunschweig (Germany), respectively.

### Description of *Stenotrophomonas muris* sp. nov

*Stenotrophomonas muris* (mu’ris L. gen. n. *muris* of a mouse). The isolate shared highest 16S rRNA gene sequence similarities to species within the genus *Stenotrophomonas* (*Stenotrophomonas maltophilia* and *Stenotrophomonas pavanii*, 99.72%). GTDB-Tk classified the genome under the genus *Stenotrophomonas* as ‘Stenotrophomonas maltophilia_F’. The highest POCP value was to the genome of *S. maltophilia* (87.40%), which also supports classification within the genus *Stenotrophomonas*. The genome tree placed the genome in the same clade as *S. maltophilia, S. pavanii*, and *Pseudomonas geniculate* (synonym: *Stenotrophomonas geniculate*). However, none of these species shared ANI and GGDC value above 95% and 70%, respectively, with the genome of this isolate, confirming its status as a novel species. In total, 261 CAZymes were identified within the genome, along with the pathway for starch utilisation. KEGG analysis identified pathways for L-glutamate production from ammonia via L-glutamine (EC:6.3.1.2, 1.4.1.-). The following antibiotic resistance genes were detected: resistance-nodulation-cell division (RND) antibiotic efflux pump (ARO:0010004), *aph(3’)* (ARO:3000126), and L1 family beta-lactamase (ARO:3004215). The 16S rRNA gene sequence of the species was most prevalent in the rhizosphere (53.1% of 1,000 samples positive), followed by plant microbiota (43.5%), and wastewater (40.3%). The type strain is **pT2-440Y**^**T**^ **(=DSM28631**^**T**^). Its G+C content of genomic DNA is 66.7 mol%. It was isolated from the caecal content of a TNF^deltaARE/+^ mouse.^12^

### Description of *Streptococcus caecimuris* sp. nov

*Streptococcus caecimuris* (cae.ci.mu’ris. L. neut. adj. *caecum*, caecum; L. gen. masc./fem. n. *muris*, of a mouse; N.L. gen. n. *caecimuris*, from the caecum of a mouse). Based on 16S rRNA gene sequence analysis, the isolate is considered to belong to the species *Streptococcus parasanguinis* (99.18% identity). However, GTDB-Tk classified the genome as ‘Streptococcus parasanguinis_B’. The isolate has POCP values >50 % to multiple *Streptococcus* species, including *S. parasanguinis* (89.4%, the highest value), and *S. pyogenes* (the type species of this genus, 54.6%). Genome tree analysis confirmed the genus assignment by placing the isolate within the monophyletic cluster of *Streptococcus* species. ANI values <95 % to *Streptococcus* spp. with a valid name (highest to *S. parasanguinis*, ANI: 94.8%, GGDC: 55.90%) support the proposal to create a novel species. The bacterium grows in BHI medium under anaerobic conditions, with visible turbidity observed within 2-3 days. The total number of CAZymes identified in the genome was 136. Further genome analyses predicted the ability to utilise glucose, cellobiose, and starch. KEGG analysis identified the pathways for production of acetate from acetyl-CoA (EC:2.3.1.8, 2.7.2.1), propionate from propanoyl-CoA (EC:2.3.1.8, 2.7.2.1), L-cysteine and acetate from sulfide and L-serine (EC:2.3.1.30, 2.5.1.47), and folate (vitamin B9) from 7,8-dihydrofolate (EC:1.5.1.3). The detection of the genes for ATP-binding cassette (ABC) antibiotic efflux pump (ARO:0010001) may indicate antibiotic resistance. The 16S rRNA gene sequence of the species was most prevalent in the human oral cavity (91.3% of 1,000 samples positive, at an average relative abundance of 9.11%), followed by human lung (81.3%, at an average relative abundance of 9.10%), and human gut (81.2%, at an average relative abundance of 1.74%). The type strain is **CLA-AV-18**^**T**^ **(=DSM 110150**^**T**^). Its G+C content of genomic DNA is 42.1 mol%, similar to *S. parasanguinis* (41.7 mol%). It was isolated from the caecal content of an SPF mouse.

### Description of *Terrisporobacter muris* sp. nov

*Terrisporobacter muris* (mu’ris L. gen. n. *muris* of a mouse). The closest 16S rRNA gene sequence similarity was to *Terrisporobacter mayombei* (99.23%) and *Terrisporobacter glycolicus* (99.16%). GTDB-Tk classified the genome within the genus *Terrisporobacter*. The highest POCP value of the genome was to *T. mayombei* (87.45%) and *T. glycolicus* (87.18%) and the genome tree analysis placed the isolate next to *T. glycolicus*. These analyses confirm the placement of the isolate within the genus *Terrisporobacter*. However, the ANI and GGDC values to the two *Terrisporobacter* species aforementioned (87.45%/34.60% and 87.18%/34.20%, respectively) were below species delineation thresholds, which justifies the proposal to create a novel species within the genus *Terrisporobacter*. Of note, the isolate was found to represent the same species as ‘Terrisporobacter othiniensis’,^89^ with ANI and GGDC values of 96.40% and 89.16%, respectively. However, this name has never been validated. The number of CAZymes identified in the genome was 198. Genome analysis predicted the ability to utilise glucose, arbutin, salicin, cellobiose, maltose, and starch as carbon source. KEGG analysis identified pathways for the production of acetate from acetyl-CoA (EC:2.3.1.8, 2.7.2.1), butyrate from butanoyl-CoA (EC:2.8.3.8), propionate from propanoyl-CoA (EC:2.3.1.8, 2.7.2.1), L-cysteine and acetate from sulfide and L-serine (EC:2.3.1.30, 2.5.1.47), L-glutamate from ammonia via L-glutamine (EC:6.3.1.2, 1.4.1.-), cobalamin (vitamin B12) from cobinamide (EC:2.5.1.17, 6.3.5.10, 6.2.1.10, 2.7.1.156), folate (vitamin B9) from 7,8-dihydrofolate (EC:1.5.1.3), and riboflavin (vitamin B2) from GTP (EC:3.5.4.25, 3.5.4.26, 1.1.1.193, 3.1.3.104, 4.1.99.12, 2.5.1.78, 2.5.1.9, 2.7.1.26, 2.7.7.2). The antibiotic resistance genes MFS type drug efflux (ARO:0010002) and tetracycline-resistant ribosomal protection protein (ARO:0000002) were identified. The 16S rRNA gene sequence of the species was most prevalent in pig gut microbiota (75.8% of 1,000 samples positive), followed by wastewater (54.8%), and activated sludge (53.2%). The type strain is **CCK3R4-PYG-107**^**T**^ **(=DSM29186**^**T**^). Its G+C content of genomic DNA is 28.7 mol%. It was isolated from the caecal content of an SPF mouse.^12^

### Description of *Veillonella agrestimuris* sp. nov

*Veillonella agrestimuris* (a.gres.ti.mu’ris. L. masc. adj. *agrestis*, wild; L. masc. or fem. n. *mus*, a mouse; N.L. gen. n. *agrestimuris*, of a wild mouse). The closest phylogenetic neighbours to the isolate are species within the genus *Veillonella* (max. 98.32% to *Veillonella caviae*). GTDB-Tk classified the genome as an unknown species within the genus *Veillonella*. The POCP value was >50 % to multiple *Veillonella* species, including *Veillonella parvula*, the type species of this genus (80.0%). The genome tree analysis placed the isolate within a monophyletic cluster of *Veillonella* species. ANI values <95 % to all close relatives with a valid name and to our other isolate from this genus (‘Veillonella intestinalis’; described above) support the creation of a novel species. Cells are coccoid (ca. 0.5-0.8 µm in diameter) when grown in WCA medium under anaerobic conditions for 3-4 days. The isolate appeared to have a limited CAZYme repertoire, with only 88 CAZymes identified. Genome analysis could not find any pathway for carbon source utilisation, but predicted the ability to produce L-cysteine and acetate (from sulfide and L-serine; EC:2.3.1.30, 2.5.1.47), cobalamin (vitamin B12, from cobinamide; EC:2.5.1.17, 6.3.5.10, 6.2.1.10, 2.7.1.156), and folate (from 7,8-dihydrofolate; EC:1.5.1.3). No antibiotic genes were identified. The 16S rRNA gene sequence of the species was most prevalent in the pig gut (33.7% of 1,000 samples positive), followed by human gut (20.3%), and human lung (18.3%). The type strain is **CLA-SR-113**^**T**^ **(=DSM 110088**^**T**^). Its G+C content of genomic DNA is 39.1 mol%, similar to *V. caviae* (38.4 mol%). It was isolated from the caecal content of a wild mouse.

### Description of *Veillonella intestinalis* sp. nov

*Veillonella intestinalis* (in.tes.ti.na’lis. N.L. fem adj. *intestinalis*, pertaining to the intestine). Strain CLA-AV-13 and Trib-3-CC-2-C show highest 16S rRNA gene identity values to species within the genus *Veillonella* (max. 96.64% to *Veillonella criceti*). GTDB-Tk classified the genomes as an unknown species within the genus ‘Veillonella_A’ (family *Veillonellaceae*). The POCP value of the isolates were >50% to species within the genus *Veillonella*, including *V. criceti* (the highest, 84.8-87.0%) and *Veillonella parvula*, the type species of this genus (66.7-69.2%). The genome tree analysis placed the isolates within the same clade as *Veillonella seminalis* and *Veillonella magna*. ANI values <95 % to all close relatives with a valid name, including *V. criceti* (ANI: 83.2-83.3%, GGDC: 26.9%) support the proposal to create a novel species to accommodate the isolates. Cells are coccobacilli (0.5-0.8 µm in length) when grown in BHI or WCA media under anaerobic conditions for 1-3 days. Genome analysis could not identify any genes for the utilisation of carbon sources, but detected the genes for production of acetate (from acetyl-CoA; EC:2.3.1.8, 2.7.2.1), propionate (from propanoyl-CoA; EC:2.3.1.8, 2.7.2.1), folate (from 7,8-dihydrofolate; EC:1.5.1.3), and riboflavin (vitamin B2, from GTP; EC:3.5.4.25, 3.5.4.26, 1.1.1.193, 3.1.3.104, 4.1.99.12, 2.5.1.78, 2.5.1.9, 2.7.1.26, 2.7.7.2). The detection of genes for lincosamide nucleotidyltransferase (LNU; ARO:3000221) may indicate antibiotic resistance. The 16S rRNA gene sequence of the species was most prevalent in the pig gut (19.7% of 1,000 samples positive), followed by wastewater (8.4%), and human gut (5.6%). The G+C content of genomic DNA of the species is 38.2-3.8.3 mol%, similar to *V. criceti* (38.4 mol%). The type strain is **CLA-AV-13**^**T**^ **(=DSM 110113**^**T**^). It was isolated from the caecal content of a wild mouse in Aachen, Germany. Strain Trib-3-CC-2-C (=DSM 105313) was also isolated from the caecal content of another wild mouse in Freising, Germany).

### Description of *Vermiculatibacterium* gen. nov

*Vermiculatibacterium* (Ver.mi.cu.la.ti.bac.te.ri.um. L. masc. adj. *vermiculatus*, in the form of worms; N.L. neut. n. *bacterium*, a small rod, and in biology, a bacterium; N.L. neut. n. *Vermiculatibacterium*, a worm-shaped bacterium). The closest relative to the isolate based on 16S rRNA gene sequence identity is *Flintibacter butyricus* (95.14%). GTDB-Tk classified the genome in the genus ‘Marseille-P3106’ within family *Oscillospiraceae*. Phylogenetic analysis showed the isolate as a separate branch within the cluster containing members of multiple genera within family *Oscillospiraceae* (*Oscillibacter, Intestinimonas, Flavonifractor*, and *Pseudoflavonifractor*). The highest POCP value was 50.1% to *Intestinimonas butyriciproducens* (type species of this genus), whereas values to the other type species of neighbouring genera, *Oscillibacter valericigenes* (34.1%), *Pseudoflavonifractor capillosus* (43.2%), *Flavonifractor plautii* (46.9%), and *F. butyricus* (39.7%) were all clearly below the genus delineation threshold. Based on the GTDB-Tk assignment, genomic tree analysis, and borderline POCP value to *I. butyriciproducens*, we propose to create the novel genus *Vermiculatibacterium* to accommodate this isolate. The type species is *Vermiculatibacterium agrestimuris*.

### Description of *Vermiculatibacterium agrestimuris sp*. nov

*Vermiculatibacterium agrestimuris* (a.gres.ti.mu’ris. L. masc. adj. *agrestis*, wild; L. masc. or fem. n. *mus*, a mouse; N.L. gen. n. *agrestimuris*, of a wild mouse). The species has all features of the genus. Cells grow as straight to slightly curved rods with pointy ends, looking like short worms (ca. 1.6-2.2 0 µm in length) when grown on YCFA or mGAM blood agar under anaerobic conditions for 2-5 days. In total, 100 CAZymes were identified in the genome, with only starch predicted to be used as carbon source. KEGG analysis identified pathways for the production of acetate from acetyl-CoA (EC:2.3.1.8, 2.7.2.1), butyrate from butanoyl-CoA (EC:2.8.3.8), and propionate from propanoyl-CoA (EC:2.3.1.8, 2.7.2.1). Antibiotic resistance may be conferred by the presence of tetracycline-resistant ribosomal protection protein (ARO:0000002). The 16S rRNA gene sequence of the species was most prevalent in the mouse gut (50.8% of 1,000 samples positive, mean rel. abund. 0.20%), followed by pig gut (10.5%), and bovine gut (6.2%). The type strain is **CLA-AA-M16**^**T**^ **(=DSM 112226**^**T**^). Its G+C content of genomic DNA is 60.5 mol%. It was isolated from the filtered (0.45 µm) caecal suspension of a wild mouse.

### Description of *Weizmannia agrestimuris* sp. nov

*Weizmannia agrestimuris* (a.gres.ti.mu’ris. L. masc. adj. *agrestis*, wild; L. masc. or fem. n. *mus*, a mouse; N.L. gen. n. *agrestimuris*, of a wild mouse). Based on 16S rRNA gene sequence analysis, the bacterium is considered to belong to the species *Weizmannia coagulans*, the type species of this genus (99.65% sequence identity). The highest POCP value (81.8% to *W. coagulans*) and genome tree analysis, which placed the isolate within the monophyletic cluster of *Weizmannia* species, confirmed the genus status. However, GTDB-Tk assigned the genome to the species ‘Weizmannia coagulans_A’. Moreover, the ANI and GGDC values to *W. coagulans* ATCC 7050^T^ were 94.7% and 59.4%, respectively, supporting the proposal to create a novel species. The isolate was found to be the same species as *Weizmannia coagulans* 36D1,^90^ with ANI and GGDC values of 98.3% and 85.3%, respectively. However, strain 36D1 has never been described to represent a novel species. The isolate grows on WCA medium under anaerobic conditions; visible turbidity can be observed within 3 days. Genome analysis predicted the ability to utilise arbutin, salicin, cellobiose, sucrose, trehalose, and starch. The genes for production of the following metabolites were also detected: acetate (from acetyl-CoA; EC:2.3.1.8, 2.7.2.1), propionate (from propanoyl-CoA; EC:2.3.1.8, 2.7.2.1), L-cysteine and acetate (from sulfide and L-serine; C:2.3.1.30, 2.5.1.47), L-glutamate (from ammonia via L-glutamine; EC:6.3.1.2, 1.4.1.-), folate (from 7,8-dihydrofolate; EC:1.5.1.3), and riboflavin (B2, from GTP; EC:3.5.4.25, 3.5.4.26, 1.1.1.193, 3.1.3.104, 4.1.99.12, 2.5.1.78, 2.5.1.9, 2.7.1.26, 2.7.7.2). In addition, sulfate reduction to sulfide was also predicted (EC:2.7.7.4, 2.7.1.25, 1.8.4.8, 1.8.1.2). No antibiotic resistance genes were identified. The 16S rRNA gene sequence of the species was most prevalent in the rhizosphere (20.1% of 1,000 samples positive), followed by pig gut (12.8%). The type strain is **aMCA-6-a-A**^**T**^ **(=DSM 106041**^**T**^). Its G+C content of genomic DNA is 46.7 mol%, similar to *W. coagulans* (46.9 mol%). It was isolated from the caecal content of a wild mouse.

### Description of *Xylanibacter caecicola* sp. nov

*Xylanibacter caecicola* (cae.ci’co.la. L. neut. n. *caecum*, caecum; L. masc./fem. suff. *-cola*, dweller; from L. masc./fem. n. *incola*, inhabitant, dweller; N.L. n. *caecicola*, an inhabitant of the caecum). Based on previous analyses, 16S rRNA gene sequence similarities between members of the family *Prevotellaceae* have been shown to be uninformative for the placement of novel isolates; this data is thus not included here.^87^ Phylogenomic placement of the type strain identified it as a member of the genus *Xylanibacter*, placed between *Xylanibacter rara* and *Xylanibacter oryzae*, but forming its own branch. ANI analysis to all members of *Xylanibacter*, and the neighbouring genera (*Palleniella, Leyella, Hoylesella, Segatella, Hallella*, and *Prevotella*), confirmed that the type strain represents a novel species with all values being below 90%. In the genome, 270 CAZymes were identified along with pathways for the utilisation of starch and cellulose. KEGG analysis identified pathways for the production of acetate from acetyl-CoA (EC:2.3.1.8, 2.7.2.1), propionate from propanoyl-CoA (EC:2.3.1.8, 2.7.2.1), L-glutamate from ammonia via L-glutamine (EC:6.3.1.2, 1.4.1.-), folate (vitamin B9) from 7,8-dihydrofolate (EC:1.5.1.3), and riboflavin (vitamin B2) from GTP (EC:3.5.4.25, 3.5.4.26, 1.1.1.193, 3.1.3.104, 4.1.99.12, 2.5.1.78, 2.5.1.9, 2.7.1.26, 2.7.7.2). Ecological analysis suggests that the species is most prevalent within amplicon datasets from the mouse gut (9.4%) at an average relative abundance of 1.53%. The type strain is **PCHR**^**T**^ **(=DSM 105245**^**T**^). Its G+C content of genomic DNA is 46.2 mol%. It was isolated from the caecum and colon content of a SPF mouse.

## Data availability

The 16S rRNA gene amplicon datasets generated in this work were deposited at the NCBI under experiment-specific Project IDs: ageing mice (PRJNA807268); wildling mice (PRJNA807849); OMM mice (gnotobiotic facility A in Aachen, PRJNA807946; F1 generation, PRJNA807912; gnotobiotic facility B in Hannover, PRJNA808033); cultures from filtered gut content (PRJNA812903). The near full-length 16S rRNA gene sequences and draft genomes of the isolates were deposited at the European Nucleotide Archive and are accessible under Project no. PRJEB50452. They can also be downloaded via the project data repository at https://github.com/ClavelLab/miBC.

## Acknowledgements

We are grateful to: Marzena Wyschkon (Leibniz Institute DSMZ) for processing strains for long-term conservation; Franziska Burkart, Alicia Geppert, and Anika Methner (Leibniz Institute DSMZ) for preparation of the OMM strain mixtures; Johannes Masson, Soheila Razavi, and Theresa Streidl (University Hospital of RWTH Aachen) as well as Haiying Huang (Helmholtz Center Munich) for contributing to strain isolation; Wolf-Dietrich Hardt (ETH Zurich) for providing *E. coli* strains; Ntana Kousetzi (AG Clavel, University Hospital of RWTH Aachen) and Klaus Neuhaus (TU Munich) for their help with sequencing; the DFG-funded NGS Competence Center Tübingen (INST 37/1049-1) for genome sequencing of *E. coli* strains; Andrea Leufgen (Institute of Molecular Medicine, University Hospital of RWTH Aachen) for support with mouse sampling and sample processing for flow cytometry; Sigrid Kisling and Dirk Haller (TU Munich) as well as Stefan Rosshart (University of Freiburg) for providing mouse samples. This work was performed with support by the IZKF Core Facilities Sequencing and Cytometry (Oliver Pabst) at the University Hospital of RWTH Aachen.

## Authors contributions

BS, JO, and TC initiated the project; AA, SAVJ, TCAH, TR, MB, TS, and TC planned experiments; AA, SAVJ, RdO, AP, MB, AvS, CE, RB, FH, EO-YW, EMB, NTo, and VC performed experiments; SAVJ, RdO, and MB performed animal experiments; AA, SAVJ, TCAH, RdO, AP, NTr, AvS, CE, RB, NTo, and VC analysed data; TCAH and NT performed bioinformatic analyses; AA, SAV, TCAH, TR, RdO, AP, Ntr, TS, and TC interpreted data; AA, TCAH, TR, and BA curated data; WWN, AB, RT, H-PH, FK, BS, TS, and JO gave access to essential material and infrastructure; AA, SAVJ, TCAH, and TC wrote the paper and created the figures; JO and TC secured primary funding; TC coordinated the project; all authors reviewed the manuscript and agreed with its final content.

## Figure legends

**Supplementary Figure S1:** Phylogenomic diversity and comparative features of *E. coli* strains. **(A)** Genomes of the 21 *E. coli* strains in miBC (bold letters) and those from reference strains (grey arrows) and species of neighbouring genera were used for protein-coding gene prediction using prodigal (v2.6.3)^77^ and subsequent tree calculation using PhyloPhlAn (v3.0.60)^91^ based on 400 universal marker genes at low diversity scale using RaxML (v8.2.12).^94^ The tree was visualized in iTOL (v6.5),^92^ with the scale bar depicting the average number of amino acid substitutions per site. It was rooted using the type strain of *Klebsiella aerogenes*. Genome assemblies of the miBC strains are accessible via ENA under project ID PRJEB50452. For other strains, high-quality genomes were retrieved from GTDB (Release 06-RS202)^93^ or from the ATCC (American Type Culture Collection) website, whenever accession numbers are not given in brackets. All genomes were controlled for quality using checkM (v1.0.12).^68^ **(B)** Phenotypic traits of all *E. coli* isolates (bold letters). Individual strains were tested for the presence of flagella in two manners (see methods): (i) transmission electron microscopy after negative staining (see example micrographs at the bottom of the figure); (ii) using a Flagellin Bioactivity Assay with HEK-BlueTM-hTLR5 cells. Sensitivity to phage infection was tested using spot assays and a variety of lytic phages (see methods): (i) phages T4, T7, Qbeta, and MS2 (blue), for which the reference *E. coli* strains ATCC 11303 and ATCC 23631 served as positive controls; (ii) three phages newly isolated from sewage water (grey);^20^ (iii) therapeutic phage cocktails obtained from the Eliava Phage Therapy Center, Tbilisi, Georgia (bluish green).^95^ The ability to ferment lactose was tested using EnteroPluri-Test (Liofilchem®). For all readouts, filled circles indicate positive reactions (*i*.*e*., presence of flagella, sensitivity to phages, lactose fermentation). For the HEK-cell assays, the colour gradient (light to dark) indicates the intensity of TLR5 induction (low to strong).

**Supplementary Figure S2:** Faecal microbiota of mice at different ages. **(A)** Multidimensional plot of generalized UniFrac distances (*beta*-diversity) coloured according to animal facilities and gut locations (this colour code was consistently used in all figure panels). **(B)** *Beta*-diversity throughout sampling time points for each facility and gut location pair. Respective control mice (culled at the age of 10 weeks at each the earliest and latest sampling time point) are shown in grey (light grey, earliest time point; dark grey, latest time point). Plots per gut location were scaled to the same distance allowing for direct comparison. **(C)** Richness in samples per mouse group as in panel b. **(D)** Heatmap of the prevalence and relative abundance of significant phyla, families and phylotypes identified to display time-dependent changes. The color gradient of relative abundances (from low, light grey, to high, dark grey) was scaled independently for each row (min. and max. relative abundance values are given in scare brackets next to the taxon name). Samples in which the specific taxon was not detected appear in white. Boxes indicate significant changes in the corresponding taxa and time point, the colour indicating the direction of changes overtime (red, decrease; blue, increase). Phylotypes were annotated using EZBiocloud^61^ with the closest relative with a valid name stated along with the corresponding percentage sequence identity in brackets.

**Supplementary Figure S3:** Diversity of the strains included in Oligo-Mouse microbiota models (OMM12 and OligOMM19.1). The phylogenomic tree, the occurrence of each strain in mouse gut samples, and their number of CAZymes were determined using Protologger. Branches are coloured according to phyla: Deferribacteres, pink; Bacteroidetes, blue; Verrucomicrobioa, violet; Proteobacteria, orange; Actinobacteria, green; Firmicutes, ochre. The strains included in OMM12 are written in black and red letters. The latter two species (Acutalibacter muris and Bifidobacterium animalis) showed unstable colonization of gnotobiotic mice in previous studies^31,41^ and were excluded from OMM19.1. Instead, the nine strains added to create this model are written in brown, bold letters.

**Supplementary Figure S4:** qPCR analysis of caecal content from OMM mice in facility A (Aachen). The samples were analysed as described in the methods. The number of mice in each group is indicated in the figure.

**Supplementary Figure S5:** Detailed immune phenotyping of intestinal lamina propria (LP) (SI, small intestine; Co, colon) and gut associated lymphoid tissues (MLNs, mesenteric lymph nodes; PPs, Peyer’s patches) in OMM and control mice by flow cytometry. All leukocytes were initially gated as live CD45+ cells. **(A)** Myeloid cell populations were pre-gated as CD11b+ and individually identified as CD64+ Ly6C+ MHCII-monocytes, CD64+ Ly6C-macrophages, SSC^hi^ Ly6G+ neutrophils and SSC^hi^ Ly6G-eosinophils. **(B)** Dendritic cells (DCs) were identified as CD11c+ MHCII+ CD64-B220-cells. **(C-D)** T cells were identified as TCRβ+ and subdivided based on the expression of CD4 **(C)** and CD8 **(D)**. B cells **(E)** were gated as B220+ MHCII+ in the SI, colon and MLNs and CD19+ cells in the PPs. All frequencies are expressed as a percentage of live CD45+ cells. N = 9-11 mice per colonization group. Different letters indicate values that are statistically significant between groups (Kruskall-Wallis test followed by Mann-Whitney U-test for pairwise comparisons). Numbers of mice were: (i) small intestine (SI) and colon (Co); GF, n = 11; OMM12, n = 9; OMM19.1, n = 10; SPF, n = 10; (ii) mesenteric lymph nodes (MLNs); GF, n = 10; OMM12, n = 9; OMM19.1, n = 9; SPF, n = 9; (iii) Peyer’s patches (PP); GF, n = 8; OMM12, n = 9; OMM19.1, n = 8; SPF, n = 15).

**Supplementary Table S1:** Metadata of all strains included in miBC (www.dsmz.de/miBC).

**Supplementary Table S2:** Formatted amplicon sequencing data from the culture experiments with filtered (0.45 µm) mouse caecal slurries to obtain small-sized bacteria.

