## Supplementary material for "Enhanced cultured diversity of the mouse gut microbiota enables custom-made synthetic communities": Suppl Fig S1

### Supplementary Fig. S1

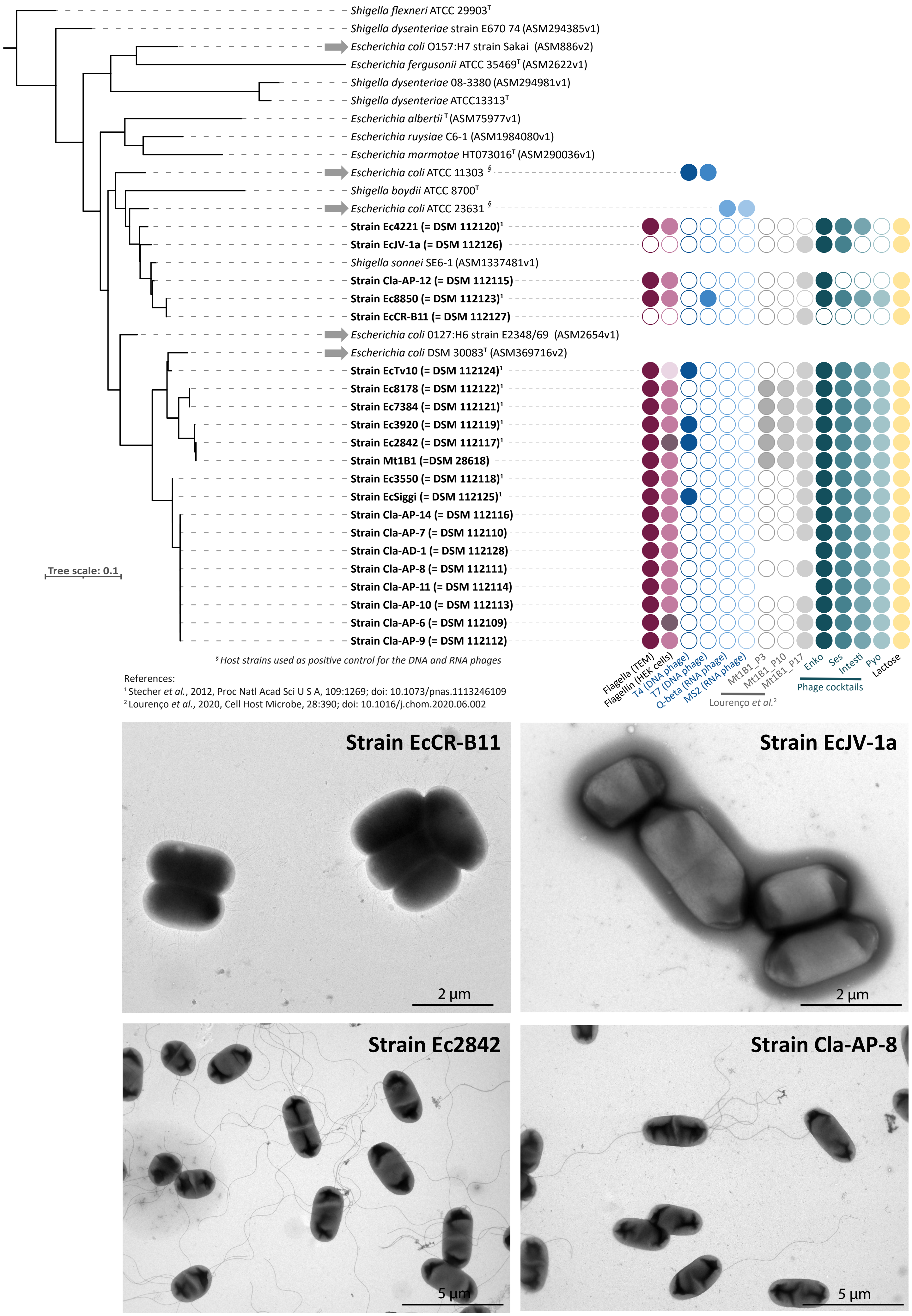
