## Supplementary figures and images for "Enhanced cultured diversity of the mouse gut microbiota enables custom-made synthetic communities"

### Suppl Fig S2

**Supplementary Fig. S2**

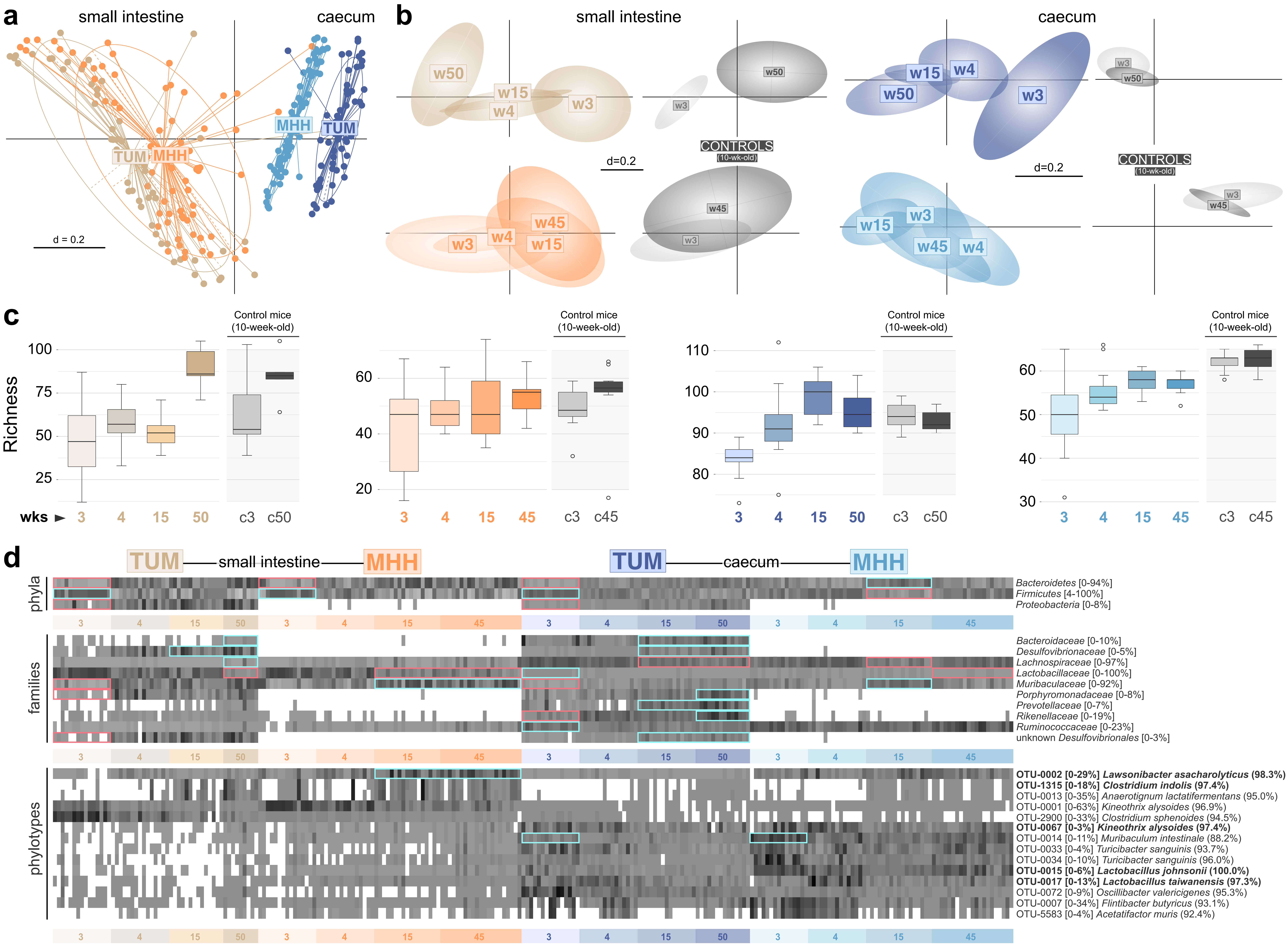

### Suppl Fig S4

#### Supplementary Fig. S4

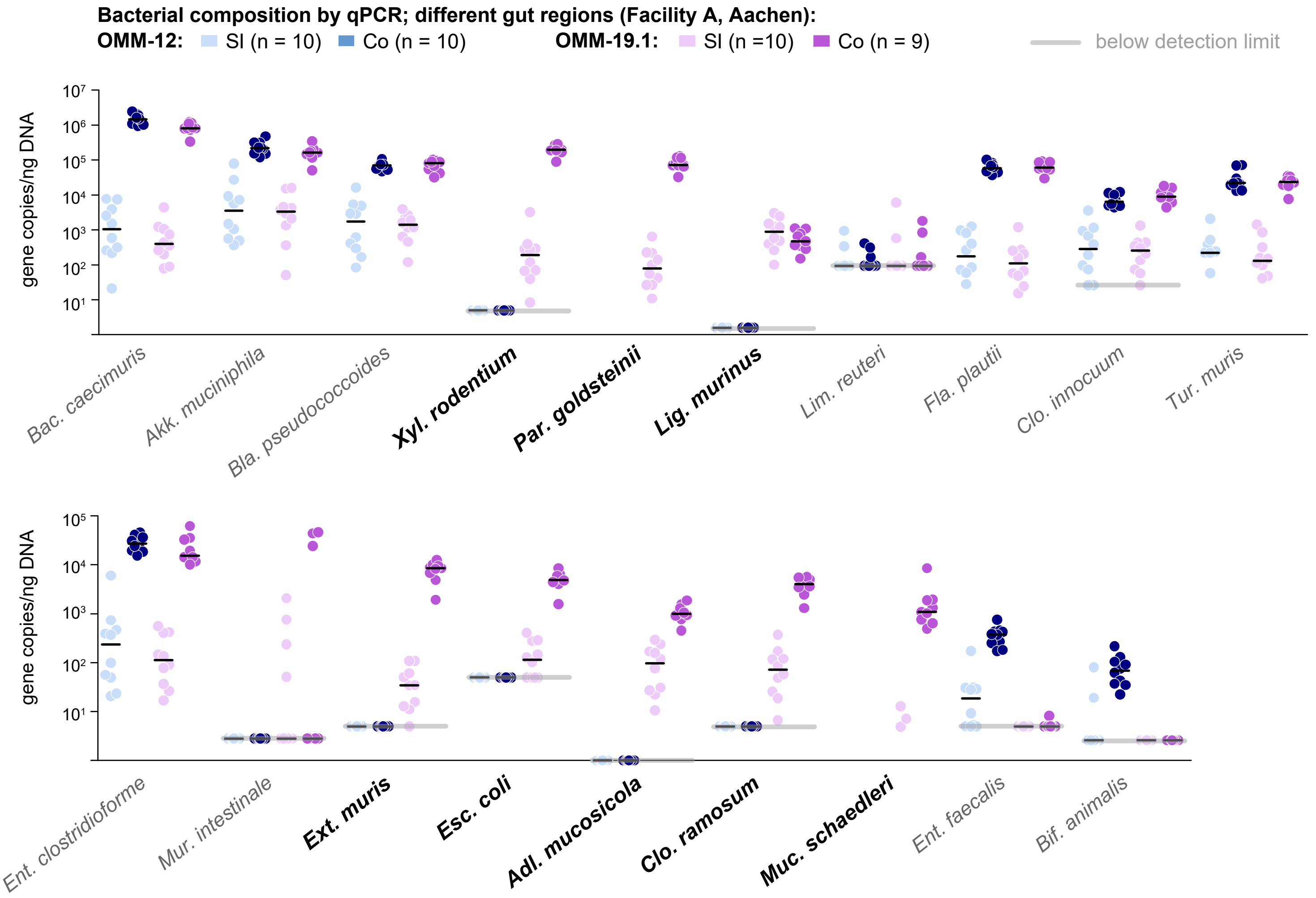

### Suppl Fig S5

#### Supplementary Fig. S5

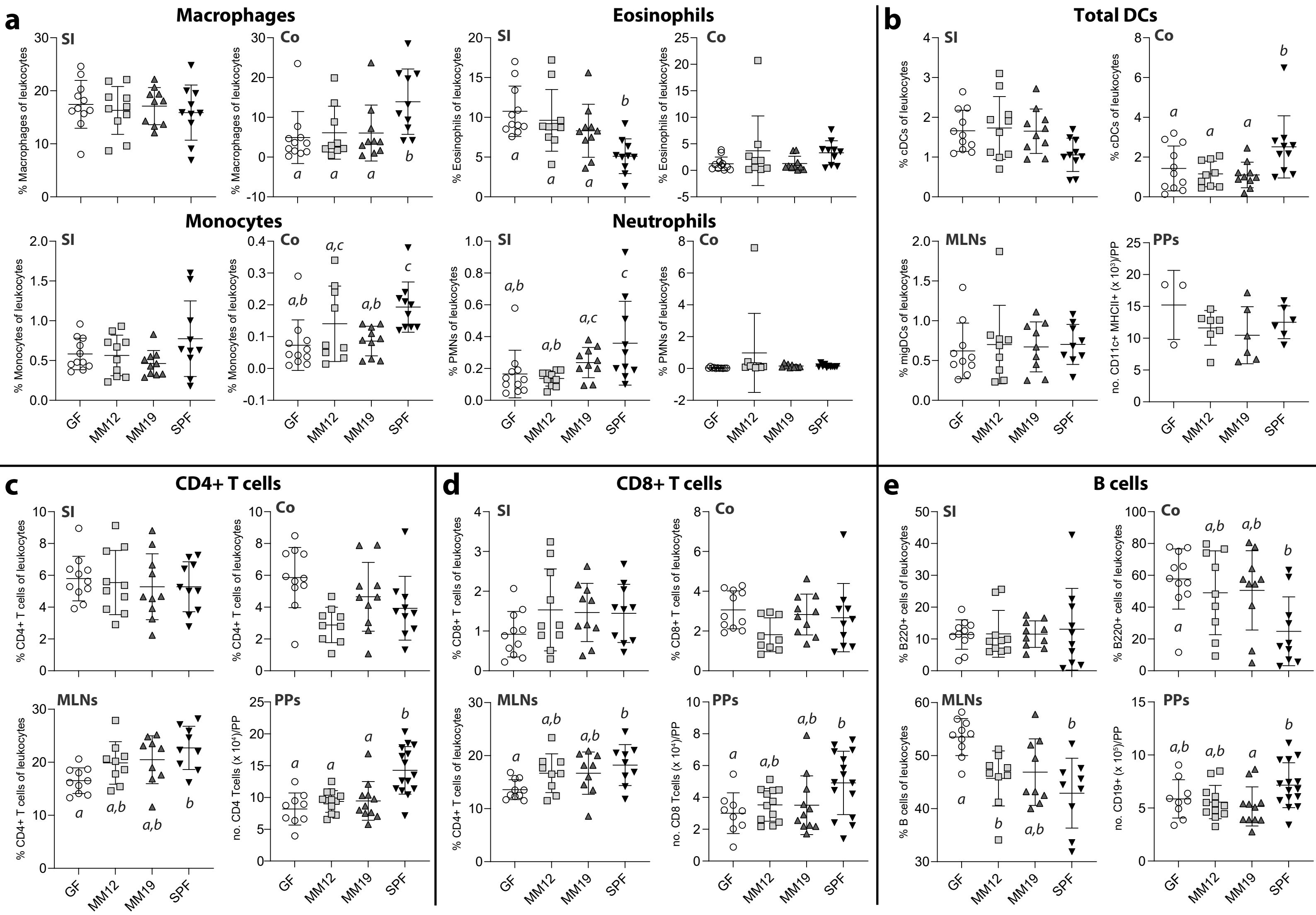
