## Supplementary material for "Enhanced cultured diversity of the mouse gut microbiota enables custom-made synthetic communities": Suppl Fig S3

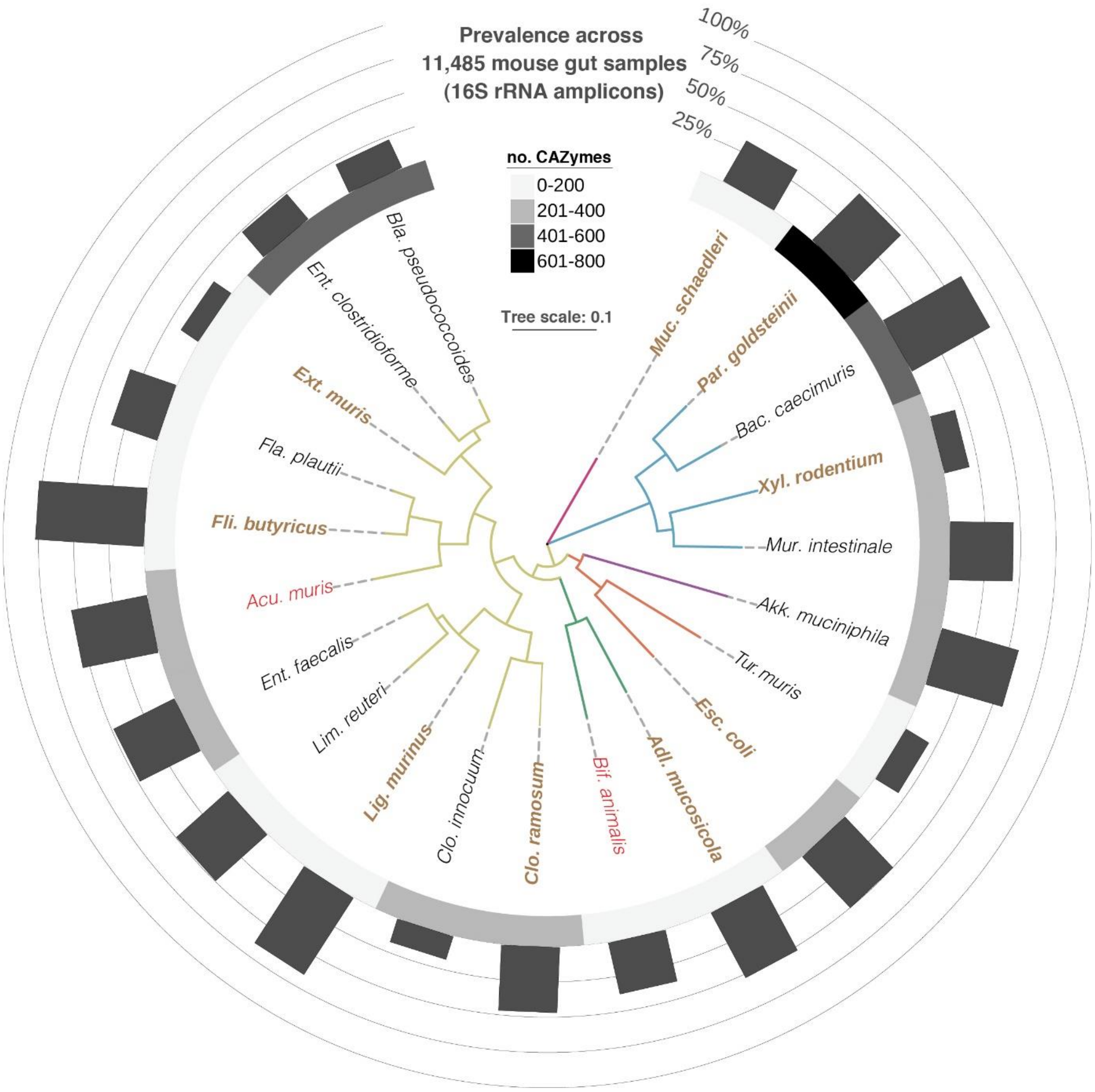


**Supplementary Figure S3**

Diversity of the strains included in Oligo-Mouse microbiota models (OMM12 and OligoMM19.1). The phylogenomic tree, the occurrence of each strain in mouse gut samples, and their number of CAZymes were determined using Protologger.[1] Branches are coloured according to phyla: Deferribacteres, pink; Bacteroidetes, blue; Verrucomicrobia, violet; Proteobacteria, orange; Actinobacteria, green; Firmicutes, yellowish. The strains included in the original OMM12 model[2] are written in black and red letters. The latter two species (*Acutalibacter muris* and *Bifidobacterium animalis*) showed unstable colonization of gnotobiotic mice in previous studies[2,3] and were excluded from OMM19.1. Instead, the nine strains added to create this model are written in brown, bold letters. Their identity and the rationale for selection (specific features) are detailed in the table on the next page.

| Species name | DSM no. | Original strain ID | Features |
|---|---|---|---|
| *Adlercreutzia mucosicola* | 19490$^T$ | Mt1B8 | Isolated from ileal mucosa;[4] Produces equol[5] and deconjugates bile acids.[6] |
| *Clostridium ramosum* | 29357 | SRB509'-5-F-B | Phylogenetic diversity (family *Erysipelotrichaceae*); suggested to play a role in metabolic health.[7-9] |
| *Escherichia coli* | 28618 | Mt1B1 | Phylogenetic diversity (phylum Proteobacteria); facultative anaerobe. |
| *Extibacter muris* | 28560$^T$ | 40cc-B-5824-ARE | Producer of secondary bile acids by 7-*alpha*-dehydroxylation.[10] |
| *Flintibacter butyricus* | 27579$^T$ | BLS21 | Prevalent species in the mouse gut; produces butyrate from amino acids.[11] |
| *Ligilactobacillus murinus* | 28683 | M-6244-3B | Enriched in mouse gut; microaerophile. |
| *Mucispirillum schaedleri* | 104751 | SH1 | Enriched in mouse gut; phylogenetic diversity (phylum Deferribacteres). |
| *Parabacteroides goldsteinii* | 29187 | BS-C3-2 | Dominant species; high number of CAZymes. |
| *Xylanibacter rodentium* | 105243$^T$ | Janvier 1a | Additional member of phylum Bacteroidetes (novel genus within family *Prevotellaceae*).[12] |
