## Supplementary material for "Enhanced cultured diversity of the mouse gut microbiota enables custom-made synthetic communities": Suppl Text

**Supplementary Results:**

The aim of this part of the work was to provide a longitudinal overview of the fecal microbiota of laboratory mice. To obtain a comprehensive dataset, samples from two animal facilities (MHH, Hannover Medical School, Institute of Laboratory Animal Science; TUM, Technical University Munich, School of Life Science Weihenstephan) were included. Mice at the age of 3, 4, 15 and 45 (MHH) or 50 (TUM) weeks had their caecum and small intestine sampled for 16S rRNA gene amplicon sequencing. To control for environmental changes within each facility, 10-week-old control mice housed at the respective site were sampled simultaneously to the experimental mice at the earliest and latest time points (ca. one year apart). After quality control, chimera-checking, and OTU filtering, 6,314,954 paired reads (averaging 17,444 ± 7,968 per sample) representing a total of 130 OTUs were studied.

The overall phylogenetic makeup of the mouse fecal microbiota was clearly separated based on gut location and the facility of origin (**Suppl. Fig. S2a**). Hence, to explore the effect of ageing, each facility-gut location pairing was studied individually. Phylogenetic distances showed successive changes within the microbiota of both gut locations from TUM and the small intestine samples from MHH (**Suppl. Fig. S2b**). The greatest separation between time points occurred between week 3 and 4, when the mice were weaned. Differences in profiles between samples at the latest time point vs. 15 weeks of age were clearly evident in TUM mice. However, less difference was observable between MHH samples over time. The control samples for each gut location-facility pair confirmed that the changes in *beta*-diversity of the ageing mice was independent of changes within the respective facility (facility-dependent overtime shifts were only observed in the small intestinal samples of TUM mice and were opposite to changes observed for ageing mice).

From week 3 to week 15, the number of detected species (richness) was shown to increase within caecum samples in both facilities. However, all small intestinal samples consistently contained approx. 50 species, apart from the week-50 samples from TUM (**Suppl. Fig. S2c**). This unique feature was accompanied by a significant increase in the diversity of bacterial species within the environment, as shown via the control samples. This may suggest that the number of bacterial species present within the murine gut is not limited by age and change in the environment of the host can lead to further expansion of the host microbial diversity.

Age-dependent changes in the relative abundance and prevalence of phyla, family and phylotypes were studied and compared against their respective controls to ensure changes were not due to environmental factors (**Suppl. Fig. S2d**). Members of the phyla *Bacteroidetes* and *Firmicutes* had opposite patterns during ageing ($R^2$ = -0.99, $p<0.0001$), with *Bacteroidetes* significantly increased between the early (week 3 and 4) to mid (week 15) and late (week 45-50) life samples, whilst *Firmicutes* significantly decreased, particularly within the small intestinal samples of mice from both facilities. Members of the phylum *Tenericutes,* which was singularly represented by the family *Anaeroplasmataceae,* increased in relative abundance from week 3 within the caecum samples of both facilities. However, this increase was only significant in the samples from MHH. Whilst present in animals from both facilities, *Proteobacteria* only showed a significant increase in relative abundance from early to late life within the small intestinal and caecal samples from TUM.

The small intestinal samples from both facilities showed a significant decrease in the relative abundance of *Lactobacillaceae* from early to late life. This pattern was also observed in the caecum but with a lower starting relative abundance. Members of the family *Muribaculaceae* increased in relative abundance within the small intestines after weaning in both facilities and in the caecal samples from TUM. A facility-independent increase in *Ruminococcaceae* was noted within the small intestinal samples between both the early and mid-life samples when compared to the late life samples.

Multiple families showed significant patterns of change within the caecal samples from TUM, which were independent from changes in the environment as determined by the controls: *Lachnospiraceae* showed a successive decrease in relative abundance across each time point. *Rikenellaceae* and *Porphyromonadaceae* displayed a pattern of successive increased relative abundance across the time points, apart from week 4 to week 15. *Desulfovibrionaceae* and another yet unknown family within the order *Desulfovibrionales* were both specific to the TUM facility and were observed to significantly increase in both the small intestine and caecum.

At the level of single molecular species, similar patterns of changes in relative abundance and prevalence to those seen at the family level were observed. OTU-0002 (most closely related to *Lawsonibacter asaccharolyticus*, a member of the family *Oscillospiraceae*) increased in relative abundance in both facilities between pre- and post-weaning sampling. OTU-1315 (*Clostridium indolis*) and OTU-0013 (*Anaerotignum* sp. within the *Lachnospiraceae*) were observed to increase in prevalence between early and mid-life samples in the small intestine and caecum samples at both facilities. The relative abundance change of both OTUs was greater within the small intestines (<20% rel. abund) than the caecum (<2% rel. abund). This was confirmed to be independent of any changes in the environment as the controls showed consistently low prevalence at both starting and end time points. OTU-0001 (*Kineothrix* sp.) and OTU-2900 (*Clostridium* sp.) decreased in relative abundance within the small intestines at both facilities between the early to mid-life samples. The most common OTU-specific change was decreased relative abundance/prevalence within the caecal samples as 7 OTUs displayed this pattern of change (OTU-0014, OTU-0033, OTU-0034, OTU-0017, OTU-0072, OTU-0007 and OTU-5583).

Altogether, these findings show that, besides the effect of weaning, consistent changes in dominant members of the gut microbiota in mice up to the age of approx. one year are rare; results varies between animal facilities. Moreover, control mice of a definite age sampled at extreme time points should always be included in studies investigating the effects of ageing on the gut microbiota, in order to consider natural shifts happening within a facility over time. This comprehensive dataset represents a solid foundation to investigate cultured fractions of the mouse gut microbiota.
